# An implantable biohybrid nerve model towards synaptic deep brain stimulation

**DOI:** 10.1101/2024.05.31.596665

**Authors:** Léo Sifringer, Alex Fratzl, Blandine F. Clément, Parth Chansoria, Leah S. Mönkemöller, Jens Duru, Stephan J. Ihle, Simon Steffens, Anna Beltraminelli, Eylul Ceylan, Julian Hengsteler, Benedikt Maurer, Sean M. Weaver, Christina M. Tringides, Katarina Vulić, Srinivas Madduri, Marcy Zenobi-Wong, Botond Roska, János Vörös, Tobias Ruff

## Abstract

Restoring functional vision in blind patients lacking a healthy optic nerve requires bypassing retinal circuits, ideally, by directly stimulating the visual thalamus. However, available deep brain stimulation electrodes do not provide the resolution required for vision restoration. We developed an implantable biohybrid nerve model designed for synaptic stimulation of deep brain targets. The interface combines a stretchable stimulation array with an aligned microfluidic axon guidance system seeded with neural spheroids to facilitate the development of a 3 mm long nerve-like structure. A bioresorbable hydrogel nerve conduit was used as a bridge between the tissue and the biohybrid implant. We demonstrated stimulation of spheroids within the biohybrid structure *in vitro* and used high-density CMOS microelectrode arrays to show faithful activity conduction across the device. Finally, implantation of the biohybrid nerve onto the mouse cortex showed that neural spheroids grow axons *in vivo* and remain functionally active for more than 22 days post-implantation.

## Main

The ability to regrow sensory nerves to the original destinations in the adult mammalian brain is restricted [1] due to the growth-inhibitory environment of the extracellular matrix in the central nervous system, which prevents axonal regeneration [2, 3]. Optogenetic [4, 5, 6], electrical [7, 8], ultrasound [9] and transcranial magnetic stimulation techniques [10] have been developed to stimulate denervated brain regions. For sensory restoration applications, stimulation interfaces primarily target the cortex due to its easy accessibility. However, the scale of the neural ensembles and the complexity of processing in the cortex has hindered attempts at sensory restoration using cortex mounted electrodes [11, 12]. Stimulating the primary target regions of sensory organs could lead to a more natural perception of the information, since the higher level sensory processing performed in the cortex is maintained [13, 14]. Targeting these regions is not without challenges: (1) they are often located in deep brain regions making access difficult and (2) the dense packing of neurons necessitates high specificity for stimulation targets. [15]. Commercially available deep brain electrodes have up to 8 electrodes that each stimulate a cubic millimeter of neural tissue including thousands of neurons [16] and axons. Miniaturization of the electrodes combined with improved stimulation protocols have increased the specificity to ∼10 neurons [8]; nevertheless, electrode alignment [17], lack of cell specificity [18, 19, 20, 21], and the immune response [22] limit the feasibility of this technology for long term sensory neurorehabilitation.

Biohybrid approaches, in which living cells integrated with stimulation electrodes are grafted into the tissue, have been proposed to overcome these challenges[23, 24, 25, 26, 27, 28]. While allogeneic and xenogeneic transplantations of neurons and organoids into adult brains have functionally integrated new neurons into developed neural networks[29, 30], neural implants have yet to leverage this for improving biocompatibility, cell specificity and stimulation resolution. In this work, we present an implantable biohybrid nerve model targeting deep brain synaptic stimulation (Fig. 1 a). A polydimethylsiloxane (PDMS) axon guidance structure containing spheroids of neurons is integrated with a stretchable stimulation array to enable isolated stimulation of individual units of neurons and the formation of a more than 3 mm long artificial nerve-like fascicle. The axons are led into deep brain regions by a gelatin-based conduit. Besides demonstrating the full functionality of the biohybrid implant *in vitro*, we also show the feasibility of implantation and the survival and axonal growth of neurons in the hybrid implant *in vivo*. As an outlook, specific recommendations for future studies to achieve axonal integration and synaptic stimulation for fully functional biohybrid stimulation electrodes are discussed.

**Fig. 1:**
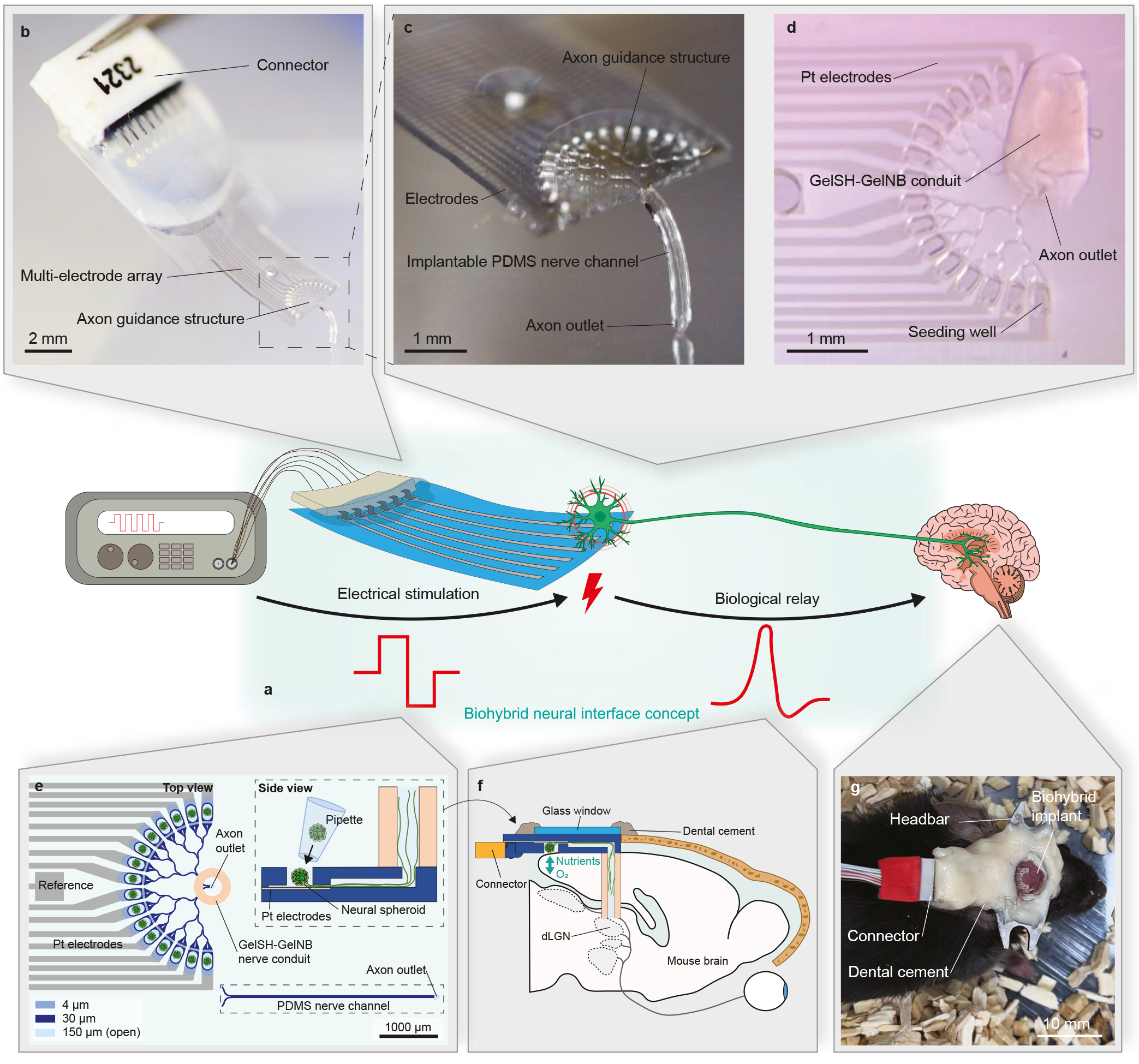
Concept and implementation of the implantable biohybrid nerve model. a: The implantable biohybrid nerve model with a PDMS based nerve channel. b: Magnification of the biohybrid axon guidance structure that is implanted on the cortex. The flexible PDMS nerve channel can be implanted into the brain to target the visual thalamus. c: Biohybrid implant in which the PDMS channel has been replaced with a GelSH-GelNB hydrogel conduit. d: Schematic of the biohybrid implant. The microfluidic axon guidance structure is aligned on top of the Pt-based microelectrode array and seeded with neural spheroids. The implantable nerve channel is made of PDMS with an axon outlet at the end or consists of a GelSH-GelNB based hydrogel conduit vertically fabricated on top of the axon outlet. Neural spheroids are seeded into each of the 16 seeding wells to grow axons towards the nerve channel. e: The biohybrid implant is flipped around and implanted onto the cortical surface of a craniotomized mouse brain. The PDMS or hydrogel nerve channel is directed towards the lateral geniculate nucleus (LGN). f: The implant is fixed onto the mouse head using a glass window and dental cement. A headbar enables subsequent mounting on a 2-Photon microscope for imaging. During stimulation experiments the implant is connected through an omnetics connector.

## 1 Results

### 1.1 Concept of an implantable biohybrid nerve model

The biohybrid device consists of a stretchable PDMS-based platinum multielectrode array, a plasma bonded axon guidance microstructure, and an omnetics connector for interfacing with external electronics (Fig.1a). The guidance structure contains 16 isolated neural seeding wells enabling stimulation of individual neural units while converging all axons into a common nerve-forming structure that can be implanted and directed towards the targeted deep brain region. We developed two implantable biohybrid nerve models: In the first model, a 3 mm long PDMS nerve channel enables the formation of the artificial nerve-like structure (Fig.1b,d). In the second model, a UV-crosslinkable and bioresorbable GelSH-GelNB hydrogel conduit replaces the PDMS nerve channel for increased biocompatibility to converge and direct axons towards their target region (Fig.1c,d).

Prior to implantation, neural spheroids are placed in each of the 16, 150 µm deep, seeding wells (Fig.1d). The biohybrid implant is flipped upside down onto the craniotomized cortical surface such that the neural spheroids receive nutrients and oxygen directly from the brain (Fig.1d,e). Neurons in the implant can grow axons along the microfluidic guidance system towards a target structure in the brain (Fig.1e). In our initial *in vivo* experiments the implant is fixed onto the mouse brain together with a metal headbar for 2-photon imaging. For electrical stimulation, the implant is connected to external stimulation electronics via a multichannel omnetics connector (Fig.1f).

### 1.2 Fabrication and characterization of the biohybrid MEA

The implant has 3 layers: a PDMS substrate, microstructured tracks and electrodes, and a PDMS microstructure for electrical insulation and axonal guidance (Fig.2a, Extended Data Fig.E1). The tracks consist of a stack of SiO2/Ti/Pt transferred on the PDMS substrate using template stripping transfer printing [31]. The tracks and electrodes have a specific meander shape on the micrometer scale [32] to improve mechanical compliance (Supplementary Fig.S1). The PDMS microstructure covers the electrode tracks up to the electrode sites assuring electric insulation. Seeding wells sit on top of the electrode with axon guidance channels leading to the nerve channel (Fig.2b). The microstructuring of the electrode enables functional calcium imaging of the underlying spheroids (Fig.2c).

The electrochemical performance of the electrodes was assessed in phosphate-buffered saline solution (PBS) (see Methods). Cyclic voltammetry (CV) was performed between -0.6 and +0.8 V (electrochemical window of Pt in PBS against Ag / AgCl [33]) (Fig.2d). According to electrochemical impedance spectroscopy (EIS) the microelectrodes exhibited an impedance of 8 *±* 1.9 kΩ (n = 13) at 1kHz. The current injection performance of the electrodes was evaluated by injecting cathodic-first rectangular, biphasic current pulses (300 *µ*s per phase, 100 *µ*s interphase delay). The current was then ramped up until the electrode polarization (removing the ohmic drop) surpassed the electrochemical window, which happens between the injection of 50 and 75 *µ*A (Fig.2d). This corresponds to a charge injection capacity (CIC) of 50 *±* 14 *µ*C/cm2 (n = 13). Next, we investigated the effect of seeding neural spheroids on the electrodes. We compared the electrochemical performance of *in vitro* (neurobasal media at 37 °C for 34 days) and *in vivo* (implanted into the mouse brain for 36 days) aged implants with new devices (Fig.2g- j). The number of working electrodes (impedance at 1kHz *<* 100 kΩ) remained unchanged for all aging groups.

However, the impedance of the explanted electrodes has increased significantly (Fig.2h). In addition, the charge injection capacity (CIC) surprisingly increased significantly during *in vitro* aging but showed the expected decrease after *in vivo* implantation compared to new implants (Fig.2i). Taken together, these results suggests that although the expected biofouling happens during implantation the resulting electrical properties of the electrodes remain within the suitable range for the proposed stimulation. We imaged a sample from each condition in a scanning electron microscope (SEM) to investigate the underlying cause (Fig. 2j). Some debris are visible on the surface of the electrodes aged *in vitro*. The origin of the debris is unknown, but we hypothesize that it is coming from the spheroids that were seeded on the electrodes. The post *in vivo* sample also shows contamination on the surface of the electrodes. It is unclear whether this is coming from the spheroids in the device or the host animal.

**Fig. 2:**
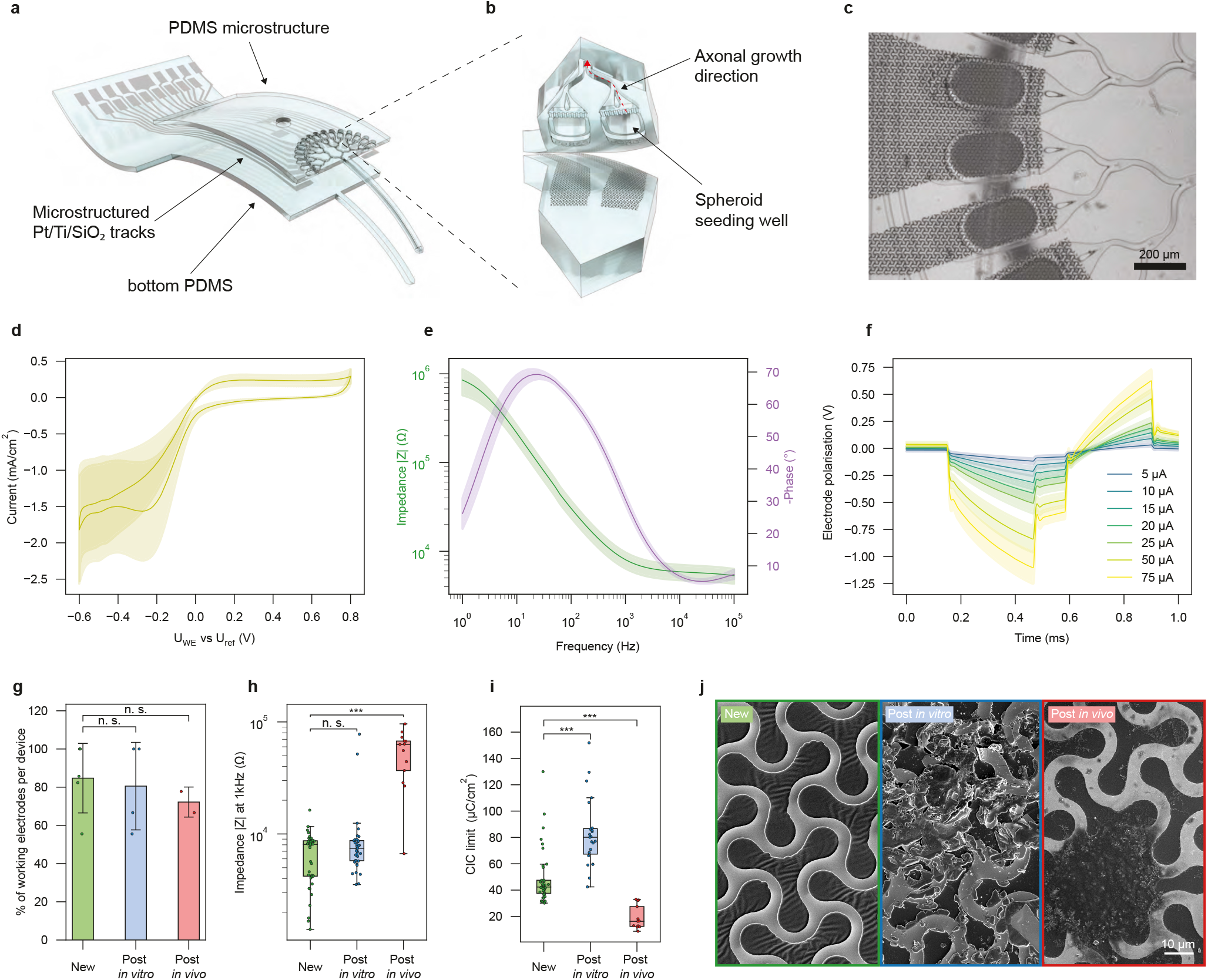
Microfabrication and electrochemical characterization of the biohybrid implant. a: Exploded view of the biohybrid device, comprised of 3 layers: a PDMS substrate, a multielectrode array, and a PDMS microstructure. b: Zoom-in on a seeding well area with the underlying microstructured electrodes. The axons of the spheroids seeded in the well area can grow along the microchannels (red arrow). c: Detailed micrograph of the electrodes, seeding wells and axonal guidance channels. d: Cyclic voltammetry (CV) of microelectrodes at 200 mV/s (n = 13 electrodes), exhibiting the electrochemical window between -0.6 and 0.8 V. e: Electrical impedance spectroscopy (EIS) of microelectrodes after microfabrication (before spheroids seeding), showing the module (green line) and phase (purple line) of the impedance versus frequency. n = 13 electrodes. f: Voltage response of microelectrodes to current-controlled biphasic pulses of 300 *µ*s per phase with a 100 *µ*s delay between phases. n = 13 electrodes, for every electrode and every current step 10 pulses were acquired and averaged. g: Comparison of the percentage of electrodes working in different conditions. New: devices after fabrication, *In vitro*: devices aged *in vitro* for 34 days. The devices were used once or twice for *in vitro* stimulation experiments. *Ex vivo*: devices measured when explanted after 36 days *in vivo*. (P = 0.16 and P = 0.5 for New vs *in vitro* and *ex vivo* respectively) h: Comparison of the electrode impedance at 1kHz in different conditions. (P = 0.16 and P = 4.4e-7 for New vs. *in vitro* and*ex vivo* respectively) i: Comparison of the charge injection capacity (CIC) of electrodes in different conditions. (P = 1.5e-7 and P = 1.4e-7 for New vs *In vitro* and *Ex vivo* respectively) j: SEM images comparing the electrode surface at different aging stages. Left: sample after fabrication, middle: sample after aging *in vitro*, right: sample after aging *in vivo*. In d, e, and f data are mean (solid lines) *±* standard deviation (shaded area). In h, data is mean (bars) and standard deviation (error bars). In boxplots, the median, quartile box and minimum and maximum values (excluding outliers) are shown. The significance was tested using a Mann-Whitney U test. The difference between two groups is considered significant if the P-value *<* 0.05, and *** denotes P *<* 0.001.

### 1.3 Axon growth within the implantable biohybrid nerve model

The biohybrid concept requires that axons (1) grow towards the nerve channel and (2) avoid turning back or growing into neighboring seeding wells. This we achieved by introducing specific geometric features into the microstructure. In timelapse movies we tracked growth cones of growing axons using a convolutional neural network and analysed how the critical parameters (*e*.*g*. channel width, number of loops redirecting axons, *etc*.) affect axonal growth speed and growth directionality (Supplementary Fig.S3,S4), resulting in the axon guidance layout shown in Fig.3a and Supplementary Fig.S5. Each of the 16 seeding wells contains 1-2 800-neuron spheroids. Neuronal spheroids extend axons through an axon filter consisting of 10×10×4 *µ*m small channels that prevent neural cell bodies from entering the channel system (Fig.3b,c and Supplementary Fig.S5c,d). The axons entering the channel systems are directionally merged (Supplementary Fig.S5e-h and Supplementary Movie 1) towards a nerve channel. At the end of the nerve channel, a triangular axon outlet facilitates axonal outgrowth for potential target innervation (Fig.3a and Supplementary Fig.S5i,j). Time-lapse recordings showed that axons enter the nerve channel in less than 16 h and reach the end of the 3.6 mm long nerve channel 44 h after seeding the retinal spheroids (Supplementary Fig.S8 and Supplementary Movie 1). Focused ion beam scanning electron microscopy (FIB-SEM) of the artificial nerve-like structure and subsequent segmentation showed that the axons form a bundle of up to 1200 axons (Supplementary Fig.S10).

**Fig. 3:**
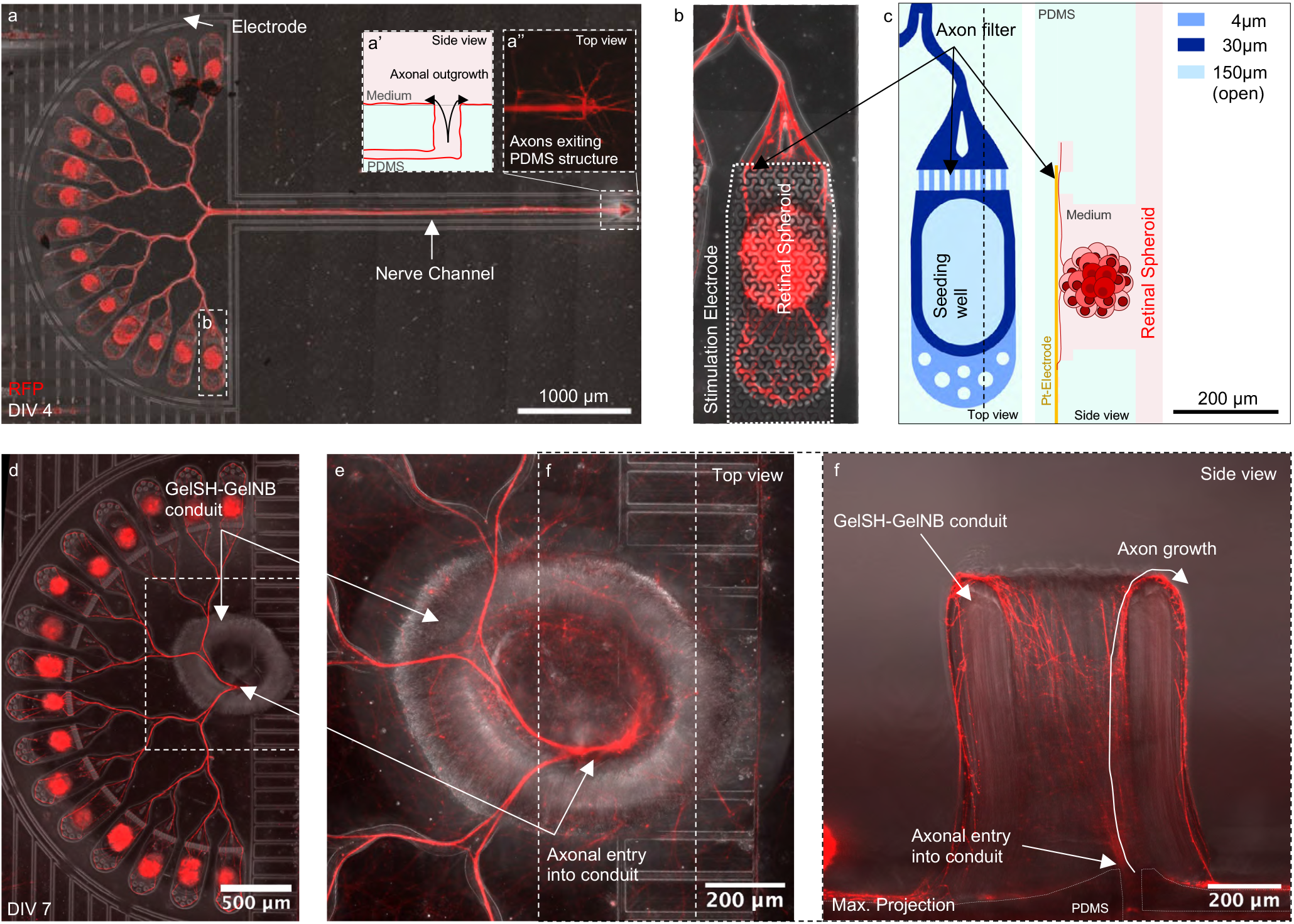
Axon growth within the biohybrid nerve model. a: Retinal spheroids grow axons through the matrigel-coated axon guidance structure to form a nerve-like structure within 3.5 days after seeding. The triangular shape favors axonal outgrowth over axon re-entry into the channel. a’: Side view of the axon outlet at the end of the nerve channel. The triangular shape favors axonal outgrowth over axon re-entry into the channel. a”: Retinal axons exit the nerve channel at the axon outlet at the end of the channel. b: One of the 16 seeding wells with an underlying electrode. Neural somata are prevented from entering the channel by an axon filter which consists of 9, 4×10 *µ*m small channels. c: Top and side view schematic of the seeding well. Retinal spheroids are manually seeded into each seeding well. d: Biohybrid implant in which the PDMS channel was replaced by a GelSH-GelNB nerve conduit directly fabricated onto the axon outlet. e: Magnification top view of the hydrogel conduit. f: Side view maximum projection of the hydrogel conduit to illustrate how axons exit the PDMS microstructure and grow up along and out of the conduit. Once they have reached the top of the conduit, the axons continued to grow along the outside walls of the conduit as there is no target structure *in vitro*.

Integration of the guided axons from the implant to the underlying tissue was achieved using a bioresorbable hydrogel [34, 35] conduit that would enable temporary guidance to the target region in the host brain followed by integration into the host tissue. Using UV-curable gel-SH gel-NB we built hydrogel conduits of about 300 µm inner diameter onto the axon outlet of the implant (Fig.4d and Supplementary Fig.S9) [36]. After 7 days, axons transited into the conduit and grew along the inside wall of the conduit (Fig.4e,f and Supplementary Fig.S13). Lateral force applied to the conduit shows that the bond to PDMS is strong enough to withstand the *in vivo* implantation procedure (Supplementary Movie 3 and Supplementary Fig.S9). Matrigel filled glass (Supplementary Fig.S11) or collagen conduits (Supplementary Fig.S12 and Supplementary Movie 2) could also serve as alternative axon-guidance structures.

### 1.4 Spike propagation within the biohybrid microstructure

Stimulation-induced action potential spikes need to reach their target and should not propagate into neighbouring stimulating channels (crosstalk). We used high-density CMOS-based MEAs (HD-MEAs) to verify that the spheroids in the guidance structures can be reliably and selectively stimulated without cross-stimulation from other source channels. For these experiments, the microstructure with the nerve-forming channel was laid flat onto the sensing area of the HD-MEA (Fig.4a). The impedance map [37] confirmed proper adhesion onto the CMOS electrodes (Extended Data Fig. E2a) and confined axonal growth within the axon guidance microstructure for up to 167 days (Fig.4b). Spontaneous activity recordings confirmed active and healthy cultures up to DIV110 (Extended Data Fig.4b). We assessed stimulation-induced spike propagation within the biohybrid structure by repeatedly (250×) stimulating the axons at each of the 16 source channels and recording the corresponding elicited action potential events within the following 3.5 ms at both the source channels and at defined locations along the nerve channel (Fig.4d, e, f). The location of the recorded spikes was color coded according to Fig.4a. We can distinguish 3 different signal propagation trends: (1) Stimulation-induced spikes propagate from the stimulated source channel towards the end of the nerve channel without entering any other source channel (no crosstalk) (Fig. 4d’ and supplementary movie 4). (2) Spikes reach the target channel but also enter into a neighbouring source channel (Fig.4e’ and supplementary movie 5) or (3) only propagate into another source channel without reaching the target channel thereby causing crosstalk between the source channels (Extended Data Fig.E2d). We stimulated the nerve at location “Target 3” to confirm results from individual channel stimulations (Fig.4a) and observed that spikes travelled into 13 out of 16 source channels (Fig.4c, f, f’ and supplementary movie 6). In summary, 7 out of 16 stimulated channels could elicit a response in the target nerve channel including crosstalk at DIV110 (Fig.4c). Stimulations of two biological replicates at DIV 27 showed a target response including crosstalk from a lower number of source channels (1/16 and 6/16) (Extended Data Fig.E2c, c’).

**Fig. 4:**
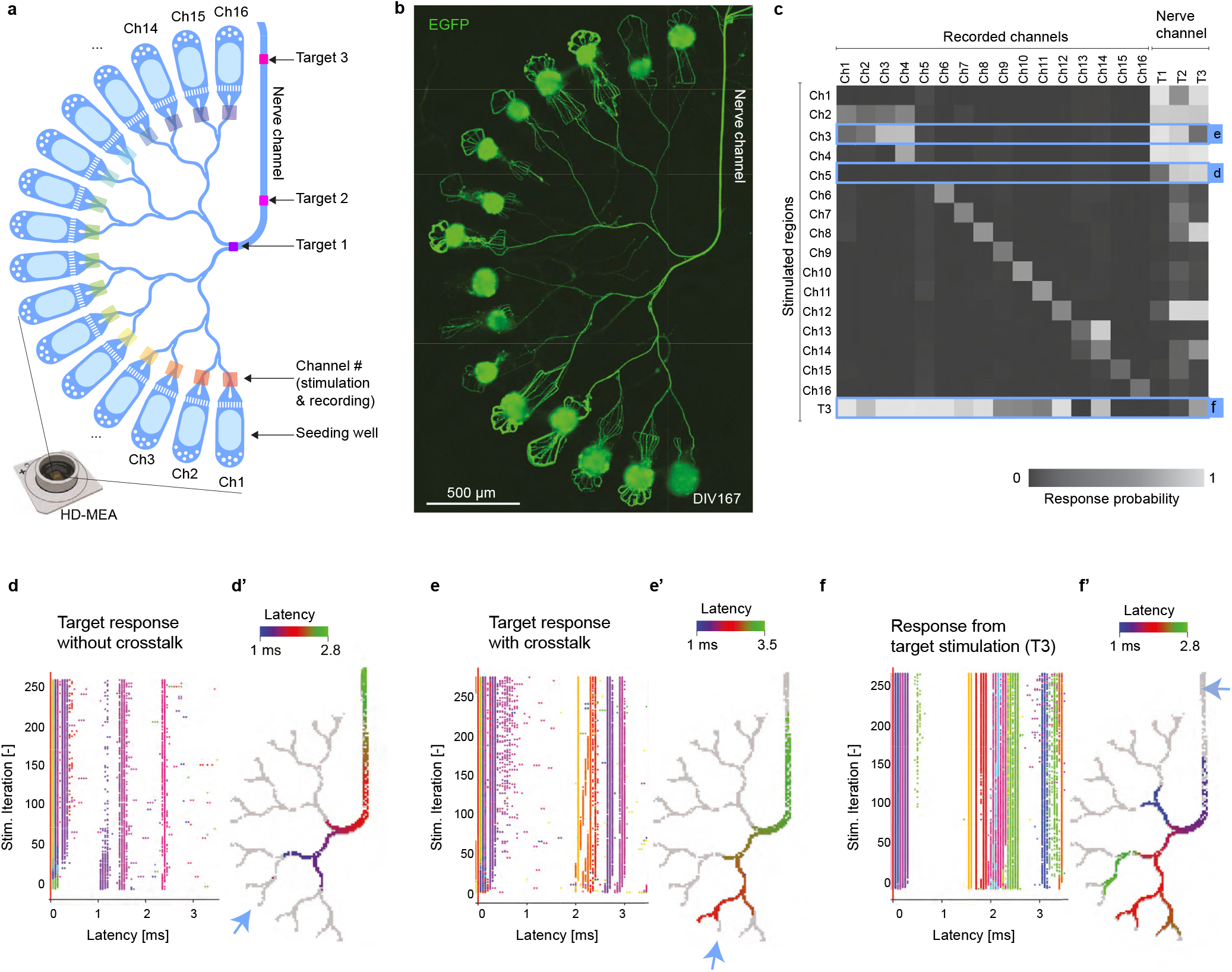
Measuring stimulation-induced action potential spikes within the biohybrid axon guidance structure to assess target response. a: Schematic of the biohybrid axon guidance structure mounted onto the HD-MEA. Colored rectangles indicate the location of specific stimulation and recording channels and regions. b: Fluorescence image showing axonal growth of retinal spheroids within axon guidance structure mounted on the HD-MEA at DIV167. c: Stimulation response matrix shows different response types (target response and crosstalk) within the biohybrid structure at DIV110. Response probability is the normalized number of detected spikes in each recorded channel or region after stimulation. (d, e, f): Stimulation induced raster plot (SIRP) of spikes propagating into the nerve channel. Colormap indicates channel location according to subpanel a. (d): response without cross talk. (e): response with cross talk. (f): Retrograde response upon target stimulation. (d’, e’, f’): Latency map outlining the elicited response propagation within 3.5ms. Blue arrow indicates stimulated channel. Grey pixels indicate all recorded electrodes. (d’): response without cross talk. (e’): response with cross talk. (f’): Retrograde response upon target stimulation.

### 1.5 *In vitro* stimulation of the implantable nerve model

Next, we stimulated the seeded spheroids on our soft, implantable microelectrode array. A PDMS well for culture media was glued around the microstructure (Fig.5a). Neural activity of spheroids in the implant (Fig.5b) was measured by functional Calcium imaging (Extended Data Fig.E3a). Electrical stimulation of working electrodes (2 V peak to peak, 200 Hz, parameters based on prior *in vitro* experiments [38]) resulted in a consistent calcium response (Fig.5c,d and Extended Data Fig.E3f,g-j). Stimulated GCaMP fluorescence propagated along the main branch (blue traces 2-4) but did not show any detectable signal in the side branches (red traces 5-6) or the neighbouring seeding wells (red traces 7-8) (Fig.5c). Neural activity could be induced in up to 50 % of the seeding wells using the tested implants (Extended Data Fig.E3).

**Fig. 5:**
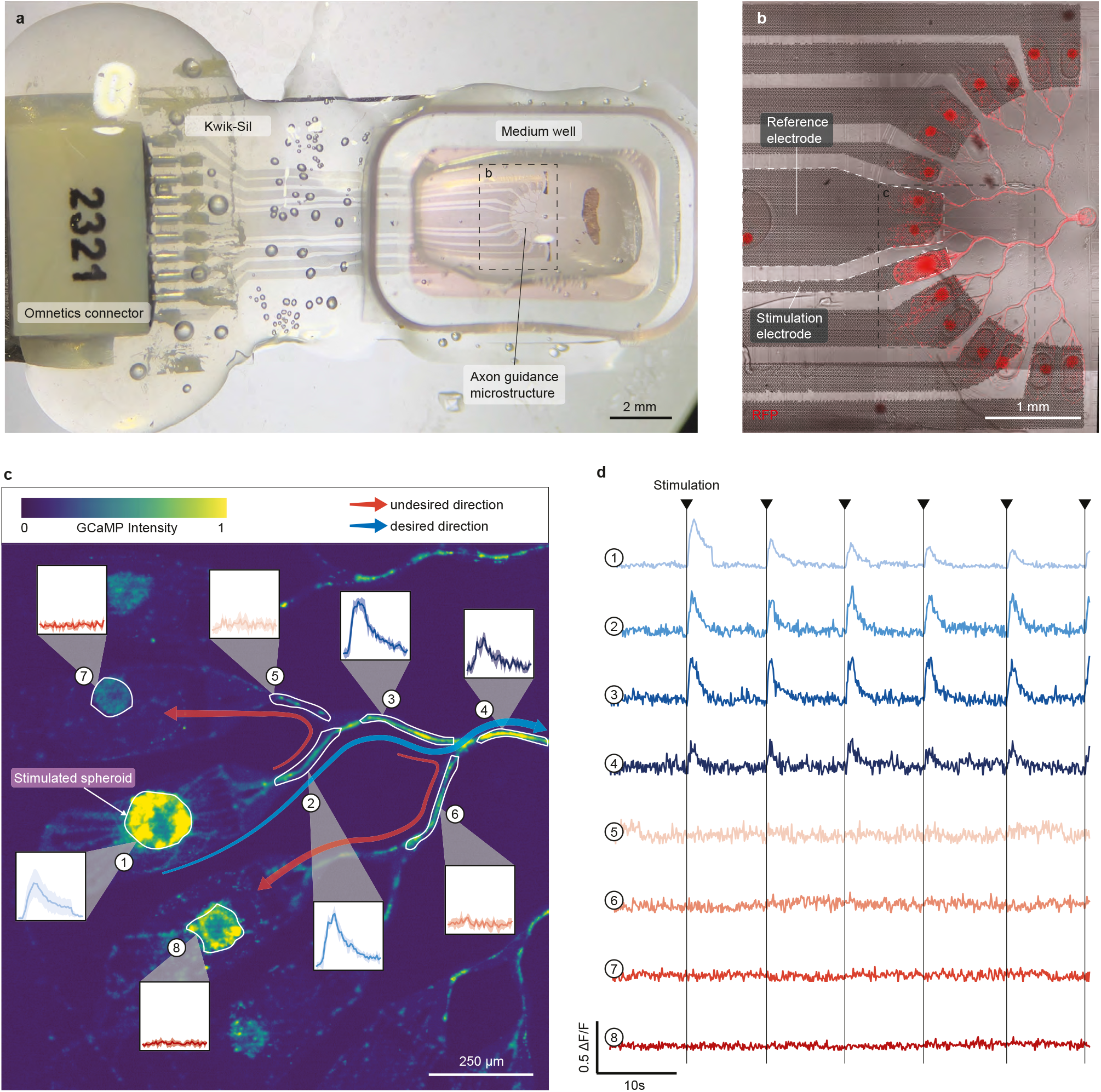
Electrical stimulation of neural spheroids in the biohybrid implant. a: Biohybrid implant with a medium well mounted on top of the axon guidance structure for *in vitro* stimulation of seeded neurons during functional calcium imaging. b: Axon guidance structure with underlying microelectrode array at DIV11. The multi-electrode array has 8 stimulation electrodes and 1 reference electrode. RFP labelled spheroids were grown within the microstructure. Image was log processed to enhance axon visibility. c: The neural spheroids were labeled with the calcium indicator GCaMP8m. One spheroid was repeatedly stimulated (2 V peak to peak, 200 Hz) and the corresponding calcium response inside the spheroid and along the channels was measured. Blue arrow indicates desired signal propagation. Red arrow indicates undesired signal propagation. d: Repeated electrical stimulation (arrows) induced a calcium response in the neural spheroid (trace 1) and along the axonal main branch (traces 2,3,4). No calcium response could be detected in neighboring spheroids (traces 7,8) or side branches (traces 5,6). Blue traces indicate traces recorded along the desired signal propagation path and red traces along undesired paths.

### 1.6 *In vivo* implantation of the biohybrid implant

The feasibility of the implantable biohybrid nerve model was tested in a living mammalian model. This involved overcoming numerous challenges due to the difficulty of fully controlling all relevant parameters within an *in vivo* environment. After a small craniotomy over the left primary visual cortex in 30-60 day old B57BL/6 mice the implant was inverted in a way that the neural spheroids were facing the brain (Fig.6a,b). In order to minimize risk of excessive bleeding, all except one device was implanted epidurally (TableS1). We prevented neural spheroids from falling out of the implant during implantation by submersing the implant in a high viscosity medium containing 1% methylcellulose. While lowering the implant manually onto the cortical surface the PDMS nerve channel was inserted towards the dorsolateral geniculate nucleus (dLGN) in the visual thalamus (Extended Data Fig.E4a). A 4-mm glass window together with a headbar were fixed on the cranium to protect the device and enable head fixation under the two-photon microscope (Fig.6a,b). Mice integrated the implant successfully with a 100% survival rate post surgery and showed no behavioral impairments in the days and weeks following the surgery (supplementary movie 9).

**Fig. 6:**
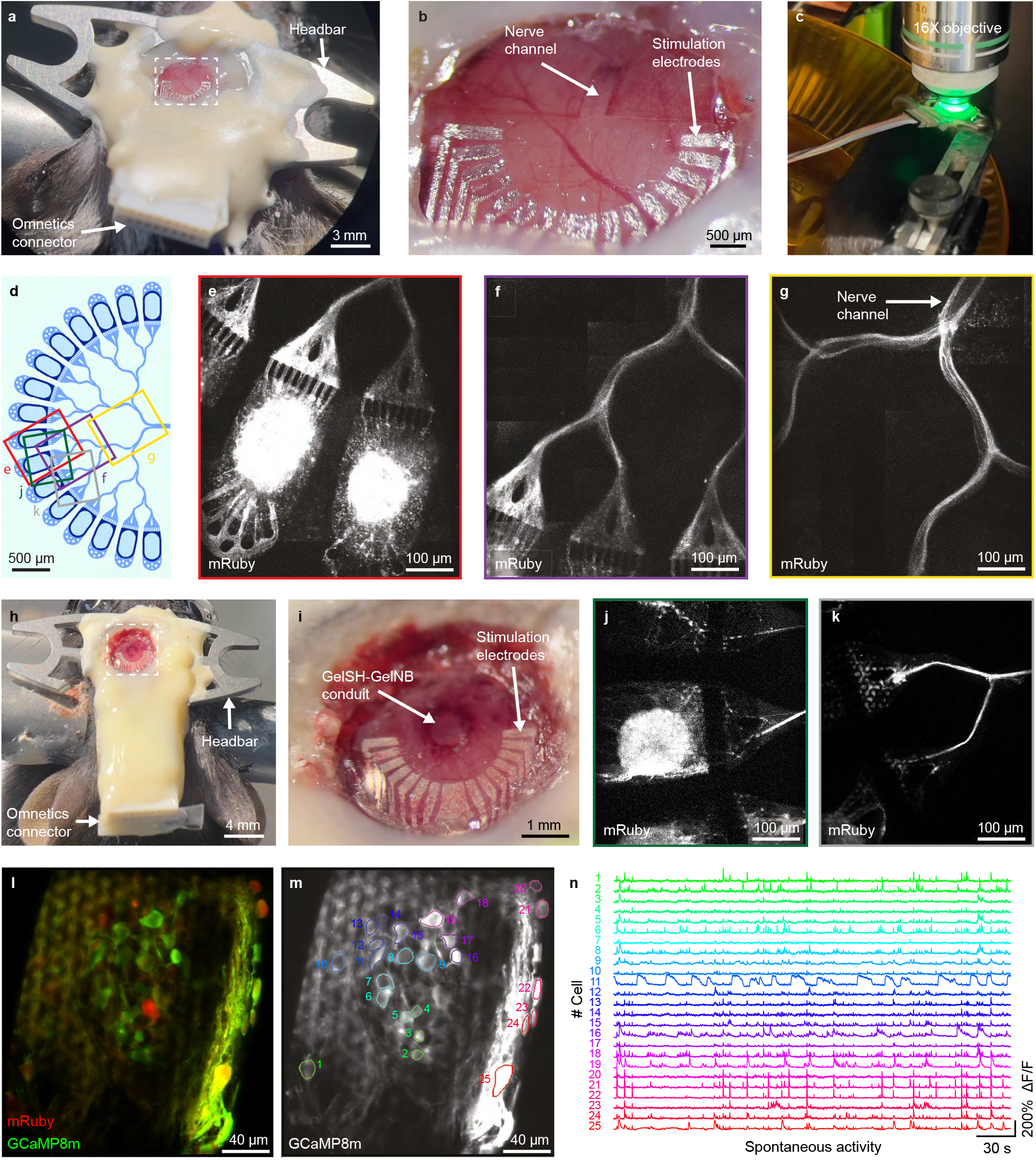
Implantation of the biohybrid implant. a: Mouse with biohybrid implant and PDMS nerve channel. b: Zoom from *a* showing the biohybrid implant on the cortex. The nerve channel enters the brain at the level of the dLGN. c: Awake two-photon imaging setup. d: Schematic layout of the implanted axon guidance structure. e: Two-photon image of cortical spheroids in the seeding wells four days after implantation. f: Two-photon image of axonal growth 4 days after implantation. g: Two-photon image of axonal growth into the nerve channel 4 days after implantation. h: Biohybrid implant with the GelSH-GelNB conduit. i: Zoomed inset from *h* showing the biohybrid implant fixed on the top of the exposed cortical surface. The GelSH-GelNB conduit enters the brain at the level of the dLGN. j: Two-photon image of cortical spheroids in the seeding wells of the GelSH-GelNB conduit four days after implantation. k: Two-photon image of axonal growth. l: Cortical spheroids expressing mRuby and GCaMP8m 22 days after implantation. m: GCaMP8m expression with ROIs for quantifications shown in n. n: Spontaneous calcium activity traces of individual neurons at 22 days after implantation.

Next, we asked how epidural implantation affects viability and axonal growth of implanted spheroids. *In vivo* two-photon fluorescence imaging (Fig.6c) showed that in DIV0 or DIV1 seeded implants, spheroids extended axons for 4-5 days after implantation in 8 out of 9 imaged mice, with axons reaching the nerve channel in 6 cases (Fig.6d- g, Extended Data Fig.E4b, Table S1). The spheroids showed intact fluorescent neural morphologies for up to 22 days after implantation (S1 and Extended Data Fig.E4c and Table S1). Devices implanted with a pre-developed 7 days old nerve structure showed intact neural structures when imaged directly after implantation (Extended Data Fig.E4d-f). For both implant strategies, axonal growth and intact axonal morphology were observed for up to 5 days (TableS1 and Extended Data Fig.E4h-i). Beyond this period, an autofluorescent (green channel) fibrous structure emerged replacing the red fluorescing axons (Extended Data Fig.E4j).

Finally, we could show that the implants with a GelSH-GelNB conduit were sufficiently robust for *in vivo* implantation (Fig.6h-i and Extended Data Fig.E4k-m). Neural spheroids showed strong mRuby expression 4 days after implantation (Fig.6k, Table S1) and axonal growth within the axon guidance structure (Fig.6k, Table S1).

Post-mortem histology showed that the GelSH-GelNB conduit integrated into the brain tissue and was still present after 22 days (Extended Data Fig.E4k-m) of implantation [34, 35].

Finally, *in vivo* spontaneous calcium (GCaMP8m) activity recordings of neural spheroids demonstrated viability and neuronal function for up to 22 days (Fig.6l-n).

Overall *in vivo* implantations demonstrate that neurons show initial healthy axonal growth down into the nerve channel and good viability within epidurally implanted biohybrid devices.

## 2. Discussion

In this work, we developed an implantable biohybrid nerve model towards synaptic deep brain stimulation. The concept is centered on electrically and functionally isolating individual neural units within an implantable nerve structure in order to exploit neurons as synaptic relays. We believe that future developments of the biohybrid strategy have the potential to increase the stimulation resolution and long term biocompatibility of currently available deep brain stimulation electrodes.

The biohybrid device was fabricated following a simple process flow. Once the template is fabricated, it can be reused several times to perform the template-stripping transfer-printing procedure, which leads to easy fabrication of MEAs on PDMS. The devices showed satisfying performance, highlighted by a successful *in vitro* spheroid stimulation as well as good electrochemical properties. The yield of working electrodes per device should be improved, but most of the non-working channels can be traced back to the interface between the omnetics connector and the Pt tracks. Future iterations might benefit from using a flexible cable connector such as that used in Fallegger *et al*. [39] to increase the number of working electrodes. When performing the aging experiment, the impedance of the electrodes after being implanted *in vivo* increases, as expected and reported in different works [33, 27, 40]. However, the impedance increase saturates at an acceptable level after 4 weeks *in vitro* with periodic stimulation. This indicates that the electrode-electrolyte interface is stable and that the electrodes do not dissolve or delaminate (a scotch tape test was performed after MEA fabrication and shows that the tracks are solidly bonded to the PDMS substrate, Supplementary Fig. S2). The CIC of the electrodes also exhibits values similar to what can be found in the literature for flat Pt [41, 42] along with the expected decrease after *in vivo* aging due to physical processes such as biomolecule absorption, fundamental differences in counter-ion transport, and diffusional limitations on charge transfer [33, 43].

We used edge guidance-based mechanisms to direct the axons into the desired direction [44]. The inherent stochastic characteristics of axonal growth introduce a level of unpredictability in directing axonal growth when only relying on-edge guidance mechanisms [45]. This may lead to signal propagation into neighboring channels and potential synaptic crosstalk. Indeed our stimulation experiments on CMOS MEAs confirmed that spikes not only propagate into the nerve channel but also into neighbouring seeding wells. It remains to be tested if the observed crosstalk can elicit secondary spikes in neighbouring channels that would negatively affect the channel selectivity and stimulation resolution. Moreover, future studies need to analyze if and how confining non-myelinated axons into an artificial nerve like structure leads to any synaptic or non-synaptic crosstalk between the axons themselves. The low number of seeding wells and the high number of neurons per seeding well in the here presented model required only four merges to converge axons from all seeding wells into the nerve channel. Therefore, the guidance motifs used in our model were sufficient to form a nerve-like structure. For future versions of the biohybrid interface, however, one should reduce the number of neurons per seeding well (ideally 1 per electrode) and increase the number of electrodes to provide useful high-resolution sensory input to the brain. Depending on the merging architecture, this will require each axon to cross a significantly higher number of channel merges, which will increase the chance of axons turning into the wrong channel. Thus, future biohybrid neural interfaces should also benefit from different guidance approaches, *e*.*g*. based on molecular gradients used to direct axons *in vivo* [46, 47].

The presented nerve model is embedded into a stretchable and soft PDMS-based stimulation array to reduce foreign body response [48] after *in vivo* implantation. The stretchability of our device was in line with what is required for our application since only rather small strain occur during implantation and on the brain once implanted. The ability to stimulate and read out functional activity may enhance existing nerve models to develop new strategies for nerve regeneration [49].

*In vivo* we have found that neural spheroids were able to grow and extend axons down into the nerve channel within the first 3 days of epidural implantation. However, about 4 days after implantation, axons started to degrade with a strong increase in autofluorescence along the axon guidance paths and seeding wells. This autofluorescence may be due to degrading neuronal structures or a triggered immune response with *e*.*g*. phagocyting macrophages or microglia [50] around the developed neuronal structures [51]. As we implanted rat neurons into wild type mouse brains, neuronal death could be the result of an immune response or because of an increased demand for oxygen and nutrients from developing neurons that cannot be matched by the passive diffusion of nutrients through the meninges. Future studies might benefit from subdural implantation into immunodeficient mice or rats, as direct transplantation of neurons, spheroids, or organoids into such models has previously led to successful neuron survival and neural network integration *in vivo* [52, 29, 30].

In summary, we have presented and characterized an implantable nerve model showing first feasibility towards *in vivo* application. Although we have yet to demonstrate axons transitioning from the implant into neural tissue to form functional synapses *in vivo* (our previous work has shown feasibility of this approach *in vitro* [38]), we present promising initial findings on neural viability and axonal growth after implantation. In future clinical applications of the biohybrid technology patient specific iPSC derived neurons can be integrated into the biohybrid device to provide personalized sensory restoration.

## 3 Methods

### 3.1 Primary neuron cultures

#### 3.1.1 Primary cell source and cell dissociation

Primary cortical and retinal cells from E18 embryos of pregnant Sprague-Dawley rats (EPIC, ETH Phenomics Center, Switzerland) were used for all experiments in compliance with 3 R regulations. Previous to the experiments, approval was obtained from Cantonal Veterinary Office Zurich, Switzerland, under license SR 31175 - ZH048/19. Briefly, E18 time-mated pregnant rats (Janvier Laboratories, France) were sacrificed, and the embryos were removed. Embryonic eyes and cortex tissue were dissected and stored in Hibernate medium on ice.

Tissues were enzymatically digested using a Papain solution (50 mg Bovine serum albumin (BSA) (A7906, Sigma-Aldrich) and 90.08 mg D-glucose (Y0001745, Sigma-Aldrich) in 50 ml sterile PBS, and vortexing 5 mL of the solution with 2.5 mg Papain (P5306, Sigma-Aldrich) and 5 µl DNAse (D5025, Sigma-Aldrich)).

The Hibernate medium was carefully aspirated from the tubes containing retinal and cortical tissue. 2.5 ml of the prepared Papain solution were added into each tube. The tubes were incubated for 15 min at 37 °C and gently shaken every 5 min. The Papain solution was aspirated without disturbing the pellets, and 5 ml warmed Neurobasal™ medium (21103049, Gibco, Thermo Fisher Scientific) with 5 % B27 supplement (17504044, Gibco, Thermo Fisher Scientific) and 10 % Fetal bovine serum (FBS) (10500056, Gibco, Thermo Fisher Scientific) was added. After 3 min of incubation, the media was removed. The incubation and aspiration steps were repeated twice using Neurobasal™ medium with 5 % B27 supplement. After the last aspiration, 2-4 ml warmed RGC medium (medium as in [53]) were added to the retina and to the cortex tissue tube. Lastly, tissues were mechanically dissociated using a 1 ml pipette until complete dissociation of the tissue.

#### 3.1.2 Spheroid generation

Commercially available AggreWell™ 400 microwell plates (AggreWell 400 24-well plate, 34415, StemCell Technologies) were coated with 500 µl of AggreWell™ Anti-adherence rinsing solution (7010, StemCell Technologies) and rinsed with 2 ml Neurobasal™ medium. The medium was replaced with 1 ml RGC (Retinal spheroids) or Neurobasal medium (Cortical spheroids) and pH-equilibrated in the incubator before seeding 960000 cells per well to achieve a spheroid size of 800 cells per spheroid. Immediately after seeding the cells were transfected with an adeno-associated virus (AAV). Retinal and cortical spheroids were transfected with a mRuby, EGFP or GCaMP8m expressing AAV (scAAV-DJ/2-hSyn1-chl-mRuby3-SV40p(A), ssAAV-DJ/2-hSyn1-jGCaMP8m-WPRE-SV40p(A), ssAAV-retro/2-hSyn1-chI-EBFP2-WPRE-SV40p(A). All adeno-associated viral vectors were provided by the Viral Vector Facility of the University of Zurich. For producing homogeneously sized spheroids, the AggreWell™ plate was centrifuged at 100 g for 3 min and transferred into the incubator at 37 °C with 5 % CO2. Spheroids were ready for seeding after 24h.

### 3.2 Biohybrid Implant Fabrication

#### 3.2.1 Fabrication of the PDMS multielectrode array

The entire process flow is illustrated in Supplementary Figure E1. The MEA tracks were first fabricated on a Si wafer and subsequently transferred on PDMS. Firstly, a 525 *µ* m thick 4 in Si wafer (Microchemicals GmbH) was patterned using standard photolithography (photoresist AZ ECI3102, spincoating 3 krpm for 30 s, softbake 60 s at 90 °C; exposure in EVG 620NT (EV Group), i-line exposure 110 mJ/cm2, post-exposure bake 60 s at 11 °C; development 60 s in AZ726 MIF, hardbake 120 s at 110 °C) and RIE etched (Oxford NGP80 (Oxford Instruments, 10 min at 100 W, 100 mTorr, 30/12/10 sccm SF6/O2/CHF3) to create the microstructured tracks layout. The wafer was then cleaned in DMSO at 80 °C for 30 min, rinsed in DI water, and briefly plasma-cleaned (3 min at 250 W, 1 mbar, Technics Plasma 100-E (Technics Plasma GmbH)). A second photolithography and etching step was carried out to delimit the individual devices (photoresist AZ P 4620, spincoating 3 krpm for 40 s, softbake 180 s at 110 °C; broadband exposure 500 mJ/cm2; development 4 min 30 s in AZ2026 MIF, hardbake 360 s at 110 °C; etching in PlasmaPro Estrelas 100 (Oxford Instruments), Bosch Process to etching depth ∼ 100 *µ*m). The wafer was then cleaned in a similar fashion as before. Prior to the material stack evaporation, the wafer was silanized (Trichloro(1H,1H,2H,2H-perfluorooctyl)silane, Sigma-Aldrich 448931), vacuum-assisted deposition) to create an anti-adhesive layer that is crucial for the subsequent transfer process. Then, a stack of SiO2/Ti/Pt (25nm/5nm/100nm) was evaporated on the wafer (E-beam evaporation, Plassys MEB550S). The MEAs were then stripped from the SI wafer and were transferred on PDMS following a procedure described in [31]. Briefly, a 15 % solution of PVA (Mowiol 18-88, Sigma-Aldrich 81365) in water was spin-coated onto a 125 *µ*m thick PEN foil (Coloprint tech-films GmbH) and cured at 60 °C for 10 min. The patterned Si wafer with the SiO2/Ti/Pt stack was heated up to 120 °C on a hotplate and the PEN/PVA was laminated onto it using a silicone roller while applying minimal pressure. At room temperature, the PEN/PVA was stripped from the wafer, effectively picking up only the patterned MEAs. The receiving substrate for the transfer was prepared by silanizing a 4 in wafer, followed by spin-coating PDMS (Sylgard 184, 1:10) at 500 rpm for 30 sec and curing it in an oven at 80 °C overnight. Once cured, the PDMS wafer and the PEN/PVA with the patterned SiO2/Ti/Pt were plasma activated (Tergeo Plasma (PIE Scientific), 35 W, duty ratio 50/255, 5 sccm O_2_ 10 sccm H_2_O, 25 s) and brought into contact to be bound together. The assembly was immediately heated up to 120 °C on a hotplate (and weighted down to ensure good contact for bonding) for at least 30 min. At room temperature, the PEN foil was peeled off the PVA. The PDMS wafer with PVA was then immersed in boiled water for 2 times 5 min with a DI water rinse in between to dissolve the PVA. The MEAs on PDMS were then cut manually using a scalpel and were carefully lifted off the support wafer for subsequent handling.

#### 3.2.2 Design of the PDMS microstructures

PDMS microstructure layouts were designed using AutoCAD (Autodesk). The layouts were sent to WunderliChips GmbH (Zurich, Switzerland) for the fabrication of a master template wafer, as well as PDMS replica that were extracted to remove uncured monomers using ethyl acetate.

#### 3.2.3 Assembly of the biohybrid implant for *in vivo* implantation

First, the PDMS MEA and microstructure were assembled using plasma bonding. Both were subjected to a plasma treatment (Tergeo, 35 W, duty ratio 50/255, 5 sccm O_2_ 10 sccm H_2_O, 40 s), then a drop of water (∼20*µ*l) was dispensed on the MEA and the microstructure was manually aligned on top using a stereomicroscope. The assembly was heated at 100 °C on a hotplate for ∼ 30 min to dry the water and ensure a good bond. An omnetics connector was aligned by hand and glued on the pads of the MEA using a silver conductive paste (Sigma-Aldrich 901769) and cured at 100 °C for 1 h. The connection points were then encapsulated in a soft silicone (Kwik-Sil, World Precision Instruments). Lastly, the device was cut to shape using a scalpel. Immediately prior coating and seeding, the whole device was plasma activated.

#### 3.2.4 Assembly of the biohybrid implant for *in vitro* stimulations

The devices for *in vitro* testing were fabricated in a similar fashion as the devices for *in vivo* implantation up to the connector gluing and encapsulating. Then, the device was glued on a glass slide and a PDMS frame was glued around the microstructure to act as a reservoir for culture medium (using Kwik-Sil as glue). Immediately prior coating and seeding, the whole device was plasma activated to increase hydrophilicity and thereby ensure that the coating solution and culture medium enter the axon guidance channel system.

#### 3.2.5 Coating and spheroid seeding for *in vivo* implantation

The implants were coated with 1:25 matrigel dissolved in RGC medium. A drop of matrigel was applied only at the nerve outlet of the microstructure to minimize surface coating and capillary forces enabled complete filling of the microstructure. The implant was then immediately submerged in RGC medium and equilibrated in a 37 °C 5 % V/V CO2 incubator.

### 3.3 FLight based gel-NB gel-SH conduit fabrication

We used thiol-norbornene clickable resins based on our previous work for the fabrication of the conduits [54]. A thiol-ene click chemistry-based system is less prone to network defects and swelling and the crosslinking chemistry is faster, when compared to chain-growth polymerization-based hydrogels such as gelatin methacryloyl (GelMA). In the present work, we used 2.5 % w/v thiolated gelatin (GelSH) and 2.5 % norbornene-functionalized gelatin (GelNB) in phosphate buffered saline (PBS). Lithium phenyl (2,4,6-trimethylbenzoyl) phosphinate (LAP) at 0.05 % w/v was added as the photoinitiator. We used the prototype FLight device developed in-house at ETH Zurich. The device is based on the filamented light fabrication technique [36], and relies on optical self-focusing and modulation instability to fabricate cm-scale constructs featuring highly aligned microfilaments. These microfilaments are prevalent throughout the length of the constructs and are excellent guidance cues for cell alignment and proliferation. For the fabrication of the devices, the PDMS devices were first plasma activated (Tergeo plasma cleaner, Oxygen, 1 min 10 s, 25 Watt), were placed in 2-well ibidi dishes (ibidi, 80286) and positioned in the Flight projection system capable of a top-down projection of filamented light. The projection images (concentric circles equaling the cross-section of the conduits) were positioned such that the open lumen of the conduits was perfectly aligned with the open channel of the PDMS devices. Next, the devices were submersed in the resin (depending on conduit length 1-2 mm), followed by a 8 s long light exposure at 64 mW/cm2 to fabricate the conduits. The constructs were then washed gently with warm PDMS at 37 °C, followed by the secondary crosslinking using transglutaminase (5 U/ml) for 30 min.

### 3.4 Microelectrodes electrochemical characterization

Electrochemical characterization of the microelectrodes was performed using a Metrohm Autolab (PGSTAT302N) potentiostat in 1x PBS. An Ag/AgCl electrode (Sigma-Aldrich Z113085) was used as reference, and a Platinum foil was used as counter electrode. The microelectrodes were characterized by performing cyclic voltammetry (CV), electrochemical impedance spectroscopy (EIS) and charge injection capacity (CIC) determination. The CV was performed between -0.6 V and 0.8 V (6 cycles, 200 mV/s). The EIS was measured between 1 Hz and 100 kHz. CV was always performed prior to EIS characterization. For the CIC determination, the electrodes were pulsed with a set current (for 300 *µ*s at negative, 100 *µ*s at 0, and 300 *µ*s at positive) 10 times. The potential of the microelectrode was recorded, and this step was repeated for increasing currents.

#### 3.4.1 Data analysis

Electrochemical characterization data were analyzed using Python 3.8 packages (Pandas, Numpy, Scipy, Lmfit, Matplotlib). Microelectrode charge injection capacity (CIC) was established by determining at what current the potential at the surface of the electrode (removing the ohmic drop) surpassed the water window (-0.6 V for cathodic pulses, and 0.8V for anodic pulses versus Ag/AgCl).

#### 3.4.2 SEM

The SEM images of the electrodes after the aging tests were taken on a TFS Magellan 400 with a beam voltage of 5 kV and current of 100 pA. Prior to imaging, the samples stored in PBS were washed with DI water and dried. They were then sputtered with PtPd (6nm) to create a conductive layer (Safematic CCU-010 HV).

### 3.5 Mechanical stability evaluation

#### 3.5.1 Adhesion test

The adhesion of Pt/Ti/SiO_2_ tracks to the PDMS was tested performing a tape test. Standard scotch tape was laminated onto the MEA, pressed down by hand to ensure good contact, then peeled off. Optical images were taken before and after to assess the robustness of the tracks.

#### 3.5.2 Stretching test

A MEA was mounted on a homemade stretch setup consisting of two linear bearings actuated by a micrometer screw each. The device was clamped onto the bearings and the assembly was brought under an optical microscope (ADD model). The device was stretched manually using the micrometer screws and pictures were taken at regular intervals.

### 3.6 Spike propagation analysis on HD-MEAs

#### 3.6.1 Mounting PDMS microstructures on HD-MEAs

HD-MEA chips were plasma activated for 1 min (air plasma) and coated with 0.1 mg/ml poly-D-lysine (PDL) in PBS for 30 min, washed three times with ultrapure water and dried using a nitrogen gun. Extracted PDMS microstructures were aligned on top of the sensing area and filled with matrigel through desiccation (1:25 in RGC medium on ice). The matrigel solution was immediately after replaced with 1 ml RGC or Neurobasal medium. Medium was changed every 3 days. Recordings were performed the day after changing the medium.

#### 3.6.2 Spheroid seeding

Primary retinal or cortical neurons were aggregated into spheroids as described in section “spheroid formation” at a size of 800 cells/spheroid and transfected with an EGFP expressing AAV (scAAV-DJ/2-hSyn1-chl-loxP-EGFP-loxP-SV40p(A)) at 15000 vg/cell. Two neural spheroids were manually seeded into each of the 16 seeding wells as described in section “spheroid generation”.

#### 3.6.3 Acquisition of spontaneous and stimulation-based electrical activity recordings with HD-MEAs

For the spike propagation analysis within the biohybrid microstructures we used HD-MEAs (26400 electrodes arranged in a grid of 220×120 electrodes, 3.85×2.1 mm2, pitch=17.5 *µ*m, MaxOne+, MaxWell Biosystems, Switzerland) with bare platinum electrodes and a flat surface topology. A maximum of 1024 stimulation electrodes can be simultaneously connected to the amplifiers for recording. We recorded from about 1000 electrodes specifically selected to cover the whole biohybrid design spanning area. Additionally, we established regions of interest at the exit of each seeding well and along the nerve channel for the stimulation-induced propagation analysis. Stimulation was performed with a single electrode. Axons inside each channel were sequentially stimulated with a rectangular cathodic-first biphasic pulse at 2 V peak-to-peak, 4 Hz and 400 *µ*s pulse width. Data were collected at a sampling frequency of 20 kHz, with a 10-bit resolution and a recording range of about ±3.2 mV, yielding a least significant bit (LSB) of 6.3 *µ*V. Using customized software developed with the MaxWell Python API, we stored the raw traces and performed spike detection with a custom algorithm [55].

#### 3.6.4 Data analysis of spontaneous and stimulation-based electrical activity recordings

Raw voltages traces were band-passed filtered (4th order acausal Butterworth filter, 300-3500 Hz). The baseline noise of the signal was characterized for each electrode using the median absolute deviation (MAD). Spikes were detected by identifying negative signal peaks below a threshold of 5 times the baseline noise. Successive events within 2 ms were discarded to avoid multiple detection of the same spike. The response upon electrical stimulation consisted in detecting spikes occurring within a time window of 3.5 ms after each stimulation pulse at each specific region of interest characterized by a set of electrodes (set of 5 electrodes at input channels Ch1-16 and set of 15 electrodes at target regions 1-3). Quantification of the response was done by binning the detected spikes with a bin size of 0.5 ms for each individual region. The response probability was calculated as the normalized number of detected spikes in each recorded channel or region after stimulation. The total expected response is the number of stimulation pulses divided by the number of selected recording electrodes at the regions). The latency maps were obtained by calculating the stimulation-triggered average spike waveform for all recorded electrodes (time window of 3.5 ms). Latency values correspond to the time of the maximum absolute amplitude of the obtained average spike waveform in this time window. For clarity, latency is only plotted for electrodes that exhibit an average spike waveform amplitude above half of the maximum waveform amplitude across all electrodes.

#### 3.6.5 Fluorescence imaging of the networks on HD-MEAs

For fluorescence imaging of the CMOS chips using an inverted confocal microscope (Olympus, FluoView 3000, Lasers: 488nm, 561nm) all excess medium was removed and a round glass coverslip (10 mm diameter) was mounted on top of the PDMS microstructure. The surface tension between the cover slip and the chip enabled us to invert the chip for imaging in an inverted microscope. For mounting in the CLSM, the CMOS chip was placed in the recording unit which in turn was mounted into a custom made metal insert that fits into the stage of the microscope. This configuration enabled inverted mounting for imaging.

### 3.7 *In vivo* experiments

#### 3.7.1 Animals

Animal experiments were performed in accordance with standard ethical guidelines (European Communities Guidelines on the Care and Use of Laboratory Animals, 86/609/EEC) and were approved by the Veterinary Department of the Canton of Basel-Stadt. For all *in vivo* experiments, both male and female 30–120 days old wild-type (WT) mice of C57BL/6 background were used, maintained on a normal 12-hour light/dark cycle, and group-housed in a pathogen-free environment with *ad libitum* access to food and drinking water.

#### 3.7.2 Preparation of the biohybrid implant for *in vivo* experiments

For *in vivo* implantation, the implants were fabricated and assembled as described in section “Biohybrid Implant Fabrication”. The implants were seeded with neurons either 7, 1 or 0 days before implantation. Implants had to be transferred from ETH Zurich to the IOB in Basel. For transport, the retinal ganglion cell medium was replaced with retinal ganglion cell medium containing 1 % methylcellulose. The resulting increased viscosity reduced medium turbulence during transport and prevented neural spheroids from being washed out of the seeding wells. For transport, a custom-made battery powered 5 % CO2 and 37 °C temperature controlled incubation box “inkugo” [56] was used. Implants were stored inside inkugo until implantation.

#### 3.7.3 Surgical procedures and implantation of the biohybrid implant

Surgeries were performed in 30-60 day old WT mice. Prior to the start of surgery, animals were anaesthetized using a Fentanyl/Medetomidine/Midazolam (FMM) mixture (0.05 mg/kg Fentanyl, 0.5 mg/kg Medetomidine, 5 mg/kg Midazolam; subcutaneous injection). Hair were removed from the head using a trimmer (Isis GT421, Aesculap). Animals were then placed in a stereotaxic frame (Model 1900, KOPF Instruments) and ocular gel (Humigel, Virbac) was applied to prevent dehydration of the cornea during surgery. A local anaesthetic (0.25 % Bupivacaine, 0.2 % Lidocaine) was injected at the site of skin incision several minutes before removing the skin using fine surgical scissors. The exposed skull was then carefully cleaned and cleared of connective tissue before being stereotaxically aligned using the anatomical bregma and lambda coordinates. A custom-made stainless steel head-mounting bar was fixed onto the skull using dental cement (Superbond C&B). After the cement hardened, a 4-mm diameter craniotomy was made over the left hemisphere with a dental drill, exposing the visual cortex and the cortical surface above the location (from skull at bregma, medial-lateral: -2.0 mm, anterior-posterior: -2.2 mm) of the dorsolateral geniculate nucleus (dLGN). Given that in most cases, devices were implanted epidurally, the dura mater was kept intact during the craniatomy. In case the device was implanted subdurally, the dura was carefully removed using a fine forceps. Using the stereotaxic setup, an injection needle (26G, 0.45 mm diameter) was then slowly inserted to a depth of 2.8 mm (from skull at bregma) to form a physical path to implant the device into the dLGN. The biohybrid device was then manually placed onto the cortex such that the neuronal seeding well openings faced the cortical surface allowing for direct nutrient support for the on-chip grown neurons. The nerve forming PDMS channel or collagen tube were inserted into the brain using the prepared physical path until the microstructure of the device aligned onto the cortical surface. A small round cover glass of 4-mm diameter (CS-4R, Warner Instruments) was placed on top of the biohybrid device and while gently applying pressure to seal the craniatomy and flatten the surface of the brain, it was carefully fixed to the skull using a cyanoacrylate adhesive (Pattex Ultra Gel). Finally, the remaining exposed parts of the device including the connector were covered in light-curing dental cement (ESPE RelyX Unicem 2, 3M) to increase their robustness for *in vivo* applications. At the end of the surgery, a Wake mix solution (0.5 mg/kg Flumazenil, 2.5 mg/kg Atipamezol; subcutaneous injection), was administered for recovery and Buprenorphine (0.1 mg/kg; subcutaneous injection) was applied to reduce postoperative pain. Analgesia was maintained by Carprofen (4 mg/kg; subcutaneous injection every 12 hours) for 48 hours.

#### 3.7.4 *In vivo* two-photon imaging of axon growth in the biohybrid implant

Axon growth in the biohybrid implant was assessed *in vivo* using a two-photon microscope (FEMTOSmart series, Femtonics). For day 0 imaging, mice were head-fixed under the microscope just after the implantation surgery, prior to the end of anaesthesia, and placed onto a heating blanket. For imaging sessions on later days, recordings were performed in awake mice, which were first habituated to head fixation on the imaging setup while being able to move on a horizontal running wheel similar to the one in their home cage. Two-photon images were acquired with a 16X water immersion objective (Nikon, NA = 0.8) and the emitted fluorescence was detected by photomultiplier tube detectors (PMTs; high sensitivity GaAsP detectors, Femtonics). mRuby was detected by a red PMT and autofluorescence was simultaneously imaged by means of a green PMT. The MESc software (Femtonics) was used to control the microscope in resonant scanning mode and the laser (InSight X3, Spectra-Physics), which was tuned to either 980 nm or 1045 nm. A transparent ultrasound gel (GEL G008, FIAB) was placed on top of the glass window in which the objective was immersed for imaging. Imaging positions in the horizontal plane were varied by moving the mouse together with head-fixation device, which was mounted on a motorized stage (MT-1078/MT-2078/MT-2278, Sutter).

#### 3.7.5 *In vivo* two-photon calcium imaging of neurons in the biohybrid implant

To record the calcium activity of neurons in the biohybrid implant, the same two-photon microscope as described in the previous section was used. The spontaneous calcium activity of the GCaMP8m+ neurons was assessed by head-fixing the mouse under the microscope while it was able to move on the horizontal running wheel. GCaMP8m was excited at a wavelength of 930 nm and frames were recorded at 26.8 Hz. Spontaneous activity in each imaging location was recorded for about 4 to 6 minutes.

#### 3.7.6 Data analysis of calcium activity recordings

Calcium activity recordings were analysed by custom-written routines in MATLAB (Mathworks). Given that recordings were performed in awake animals, horizontal motion in the imaging field was first corrected using the NoRMCorre algorithm [57]. Then, cells were manually selected on the mean projection image of the corrected calcium imaging movie using the region of interest (ROI) plugin in ImageJ (https://imagej.net). The calcium traces of a single cell were computed by taking the average fluorescence within the ROI at any given timepoint. Additionally, for each cell, to correct for non-specific neuropil fluorescence contaminating the ROI, the 30th percentile of the fluorescence within a 10-pixel wide ring at a 5-pixel distance from the ROI edge was subtracted from the mean ROI fluorescence, as it is commonly done in calcium imaging [58]. It was also ensured that the neuropil rings did not contain other cells. Finally, the calcium trace of each cell was subtracted and divided by a slow (20-s long) moving low-pass (30th percentile) filtered version of itself to obtain a normalize ΔF/F trace. The final ΔF/F trace was then filtered by a 200-ms long average filter for display purposes.

#### 3.7.7 Histology

At the end of experiments mice were deeply anaesthetized with a Ketamine/Xylazine mixture (120 mg/kg Ketamine, 16 mg/kg Xylazine; intraperitoneal injection) and transcardially perfused with cold phosphate buffered saline (PBS) followed by 4 % paraformaldehyde (PFA) in PBS. Implants were removed and brains were extracted and placed in 4 % PFA for storage at 4 °C. To confirm the correct targeting of the implant within the neural tissue, brains were washed three times for 10 min in PBS, embedded in 4 % agarose and cut at a thickness of 150 *µ*m using a vibratome (VT1000S vibratome, Leica Biosystems). Brain slices were stained with Hoechst 33342 (10 *µ*g/ml) in PBS and then embedded in ProLong Gold Antifade Mountant (Thermo Fisher Scientific). Slices were imaged around the implantation site using a confocal microscope (Olympus, FluoView 3000, Lasers: 405 nm and 561 nm).

## 4. Acknowledgements

The research was financed by ETH Zurich, the Swiss National Science Foundation (Project Nr: 165651), the Swiss Data Science Center, a FreeNovation grant, the Human Frontiers Science Program Organization, HFSPO, and an EMBO Postdoctoral Fellowship (A.F., ALT 688-2022). We thank Georg Kosche for his technical support in two-photon calcium imaging. PC would like to acknowledge funding from the Swiss National Science Foundation (Ambizione grant PZ00P2 216356; Spark grant CRSK-2 220980). The authors thank ScopeM and in particular Lucas Miriam Susanna and Stephan Handschin of ScopeM for their support and assistance in this work.

## 5 Author information

These authors contributed equally: Léo Sifringer and Alex Fratzl.

### 5.2 Contributions

T.R. conceived and designed the study. A.F. and T.R. performed *in vivo* experiments. L.S. and T.R. designed the biohybrid implants. L.S. fabricated and characterized the biohybrid implants. B.F.C. performed stimulation experiments on CMOS and analyzed spike propagation on CMOS MEAs. L.S. and T.R. performed and analyzed *in vitro* stimulation experiments. A.B. characterized axonal growth in glass tubes. T.R. characterized axonal growth in collagen conduits. S.M. provided collagen conduits. S.S. and S.J.I. optimized directionality of axonal growth. E.C., J.D., C.M.T, T.R. and L.S. optimized coating conditions for axonal growth on PDMS. L.M. and T.R. characterized axonal growth in the final implants. P.C. fabricated GelSH-GelNB conduits. L.S. and J.H. performed SEM imaging. S.J.I. designed the implant PCBs. K.V. contributed to neural spheroid preparation. J.V., B.R., S.W., S.J.I, B.M., C.M.T., K.V. and S.M. provided advice in various aspects of the project. The manuscript was jointly written by L.S., A.F., B.F.C. and T.R. and was revised by all authors.

