## Supplemental information for "An implantable biohybrid nerve model towards synaptic deep brain stimulation"

### Extended Data

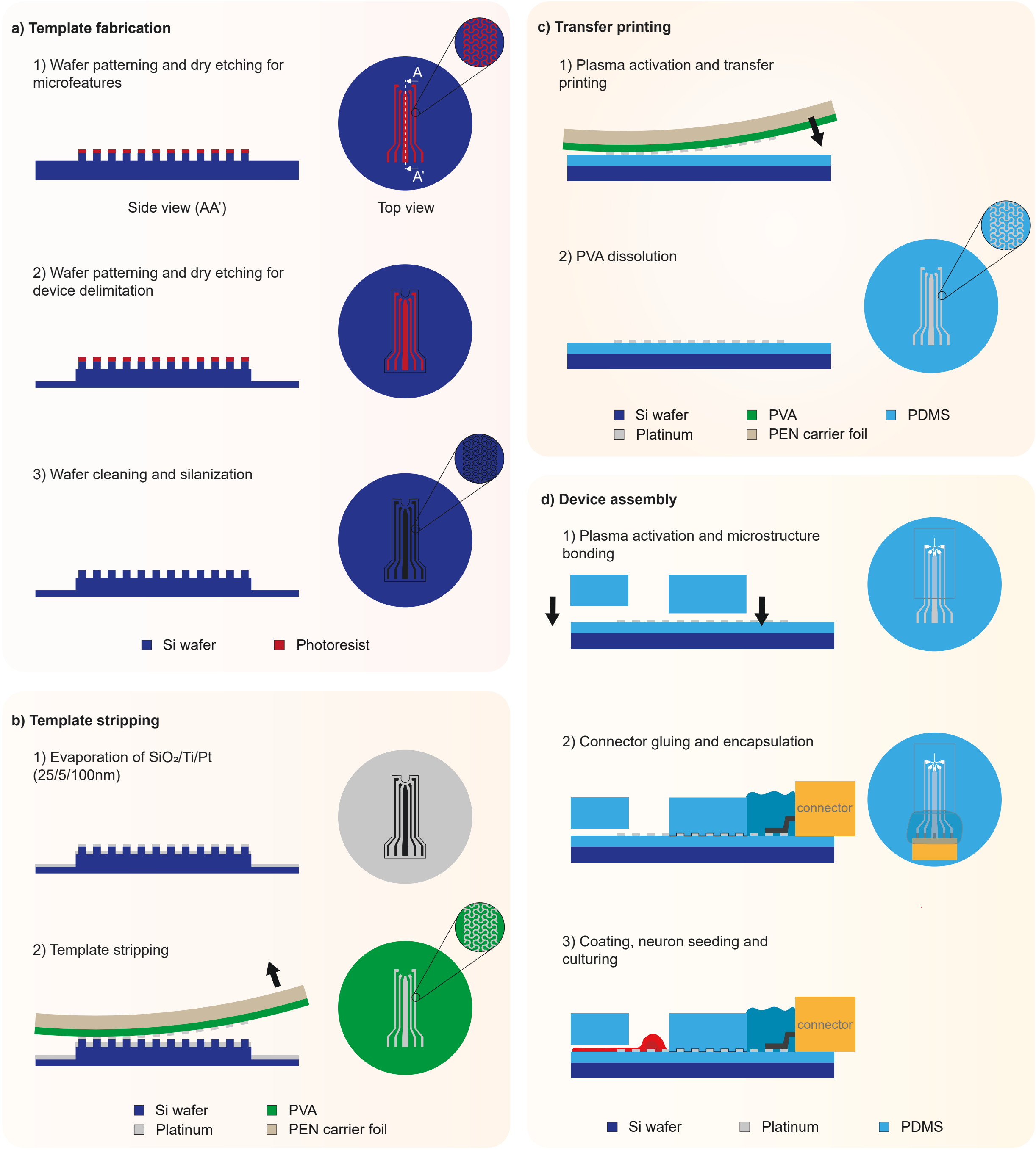


**Fig. E1**: **Fabrication process flow for biohybrid devices** a: Template fabrication: two subsequent dry etch processes are carried out to pattern the wafer; firstly to define the tracks and their micropatterning, secondly to define the device delimitation. The wafer is then cleaned and silanized. b: Template stripping: a stack of $SiO_2$/Ti/Pt (25/5/100 nm) is evaporated on the template. The template is stripped using a polyvinyl alcohol (PVA) solution spin-coated onto a polyethylene naphtalate (PEN) substrate. c: Transfer stripping: the devices are transfer printed onto PDMS (plasma activated). The PVA is dissolved in water, leaving the devices on the PDMS. d: Device assembly: the PDMS microstructure is aligned and plasma bonded onto the MEA. The connector is glued on the Pt tracks using a conductive paste. Lastly, the open connections are encapsulated with a biocompatible silicone.

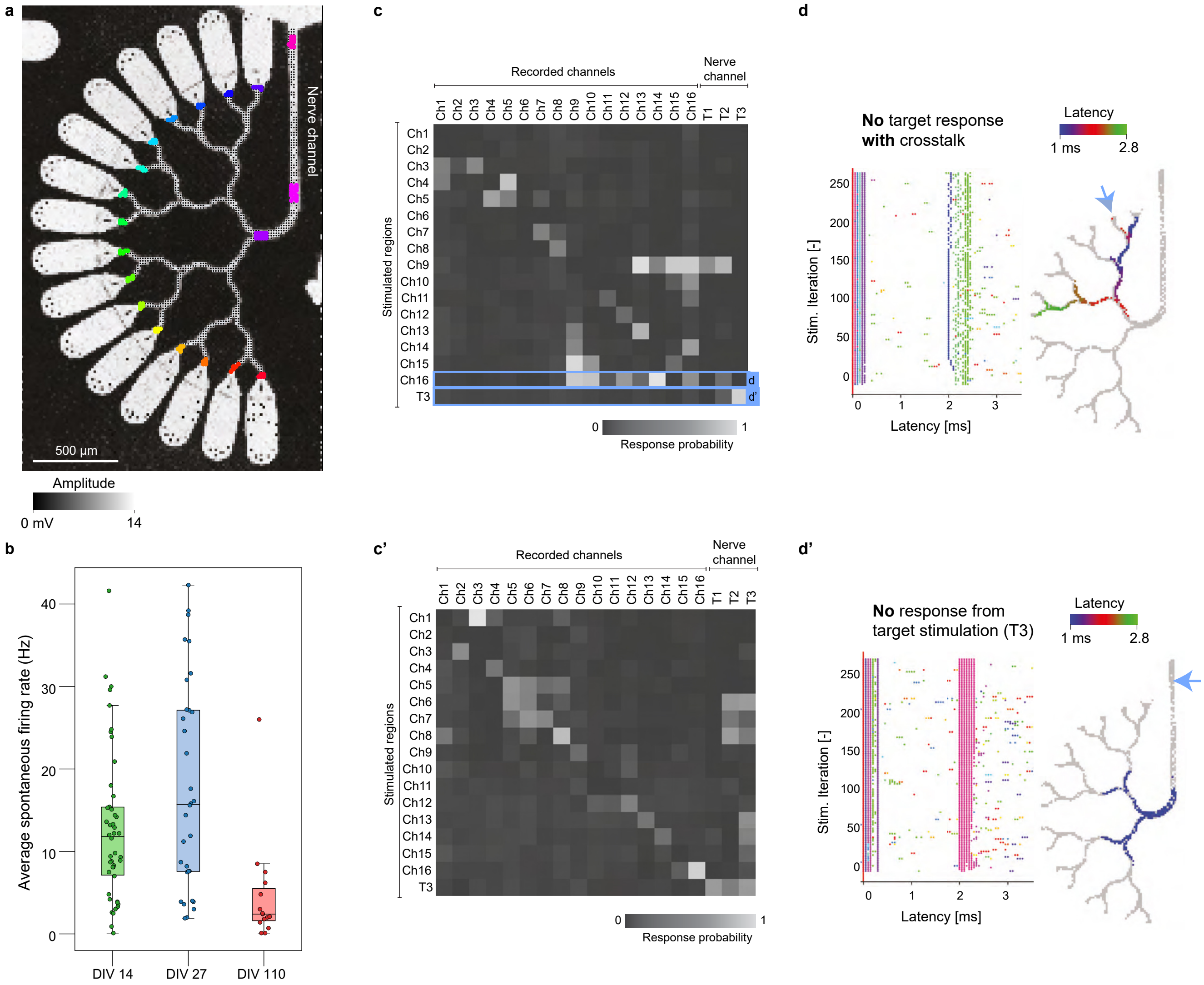


**Fig. E2**: **HD-MEA experiments to obtain response matrices of retinal neurons cultured in the biohybrid structure at DIV27** a: Impedance map of the biohybrid structure created by applying a sinusoidal voltage (14 mV peak-to-peak, 1 kHz) confirming proper adhesion of the microstructure onto the CMOS surface. Black dots within the structure represent all routed electrodes for recording. b: Box plot showing spontaneous activity. Each dot represents the average firing rate of a single channel at DIV14 (3 MEAs), DIV27 (2 MEAs) and DIV110 (1 MEA), measured at a single electrode. (c, c'): Stimulation response matrix shows different response types (target response and crosstalk) within the biohybrid structure at DIV27. Response probability is normalized number of detected spikes in each recorded channel or region after stimulation. d: Left: Stimulation-induced raster plot (SIRP) and right: Spike latency map of spikes propagating into a neighboring channel without reaching the nerve channel. Blue arrow indicates location of stimulation. d' Left: SIRP and Right: Spike latency map showing no response after stimulating the target channel.

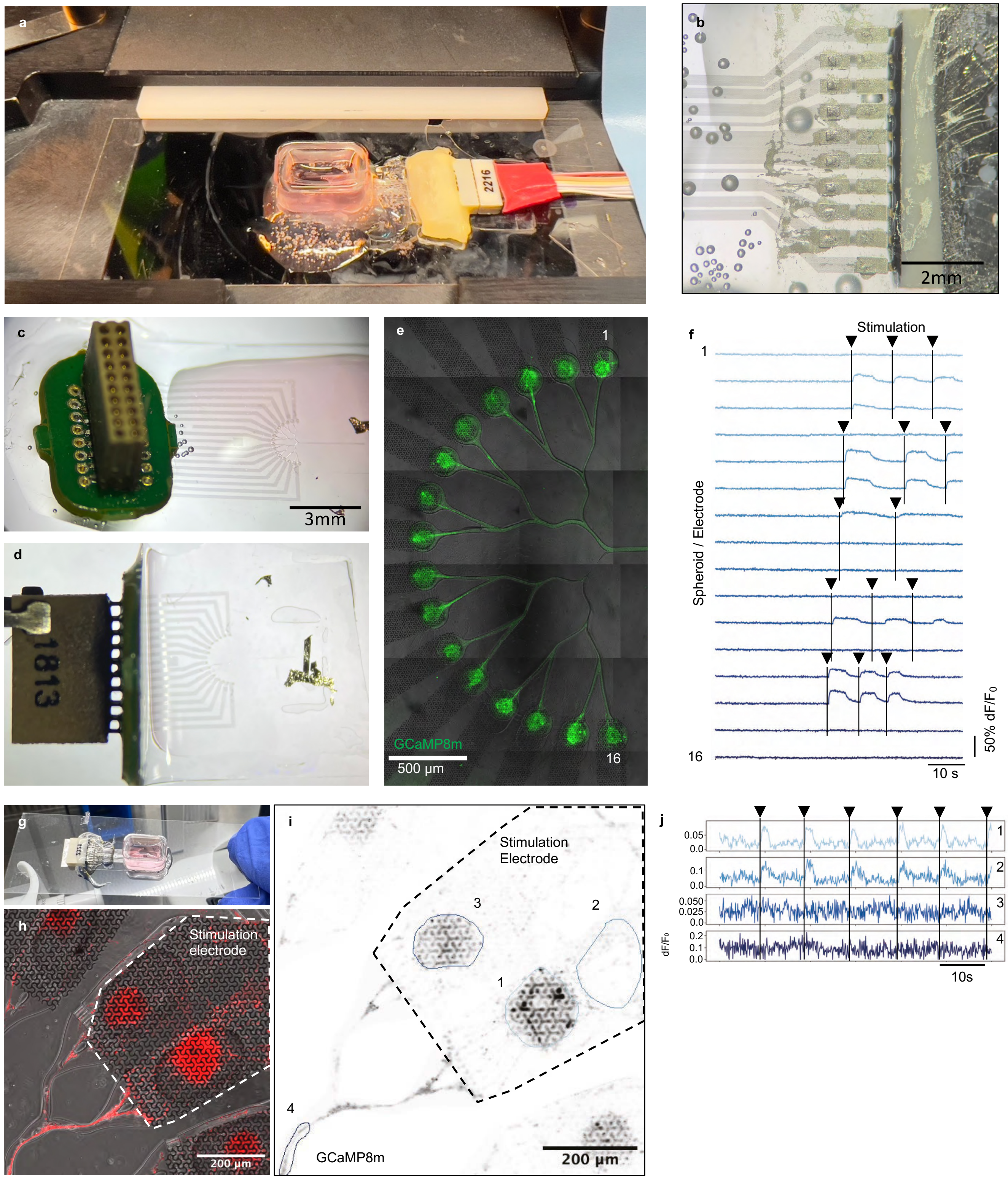


**Fig. E3**: ***In vitro* stimulation of neural spheroids inside the biohybrid implant** a: Biohybrid implant mounted on a confocal microscope headstage for functional calcium imaging. b: Bottom view of a directly mounted omnetics connector aligned with the Pt-tracks and glued with silver epoxy. c: First version of the biohybrid implant with a vertically mounted omnetics connector with a PCB in between electrodes and connector. d: Connector with PCB was bent by 90 degrees to illustrate how it would be implanted onto the cortex. e: Fluorescence image of the GCaMP8m labelled cortical spheroids. f: Neural spheroids were stimulated manually at different timepoints simultaneously (black arrows). g: Second version of the biohybrid implant with the omnetics connectors mounted directly onto the multielectrode array. The implant is mounted onto a glass slide for Calcium imaging. h: Zoom into seeding wells with underlying stimulation electrode. i: GCaMP8m fluorescence image of stimulated electrode. ROIs were manually selected to measure the Calcium response. j: Functional Calcium traces ($\Delta F/F$) in response to electrical stimulation (0.3 s, 200Hz, 2 V peak-to-peak, Inter stimulus interval (ISI): 10 s.

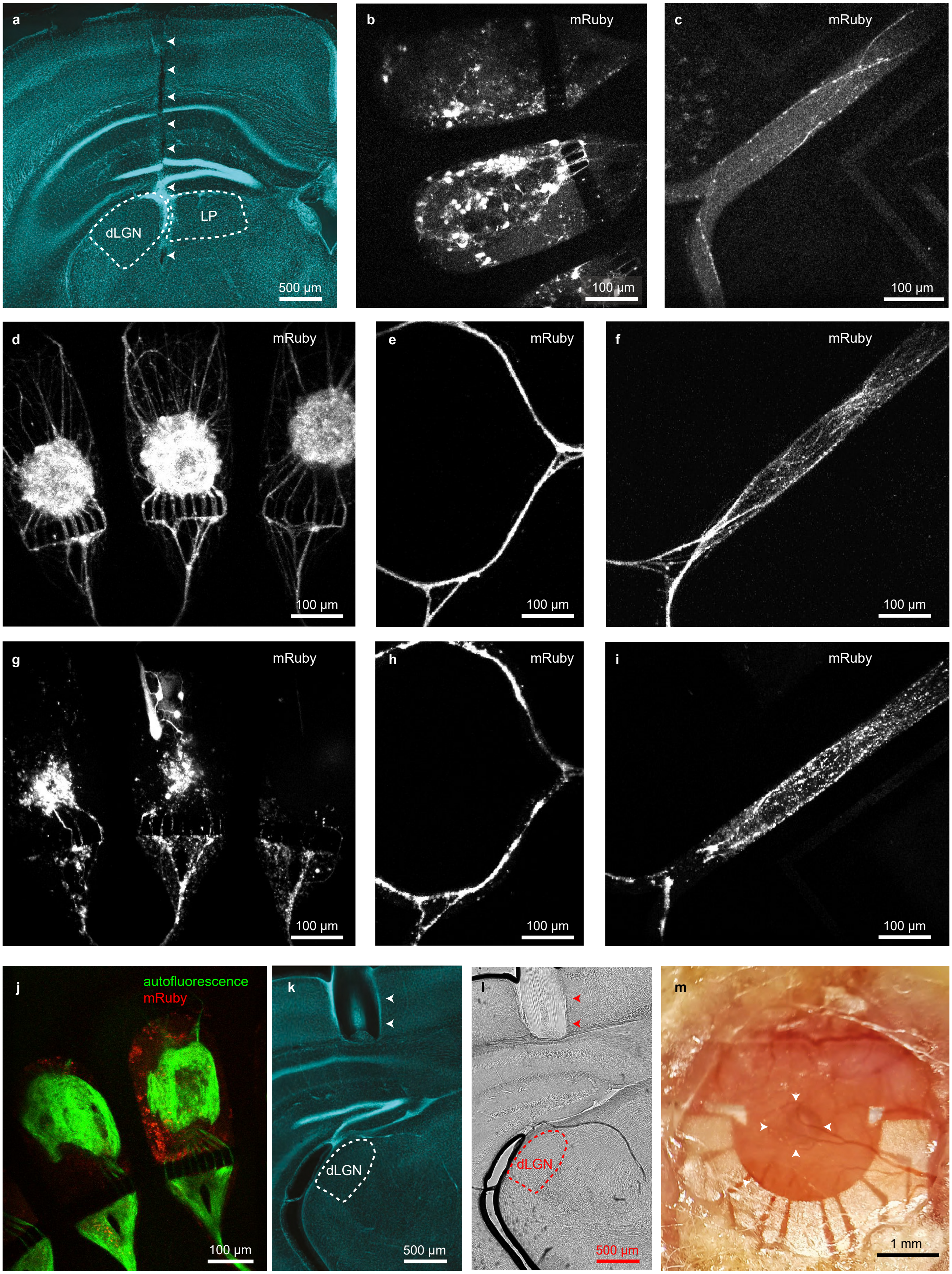


**Fig. E4**: **Properties of biohybrid implants after *in vivo* implantation** a: Implantation track of the PDMS nerve channel in the mouse brain. White arrows indicate the lesion location. Hoechst stain. dLGN: dorsolateral geniculate nucleus, LP: lateral posterior nucleus of the thalamus. b: Two-photon image of retinal spheroids in the biohybrid implant with PDMS nerve channel four days after implantation.

Figure E4 (continued): c: Two-photon image of the retinal spheroid axons. d: Two-photon image of DIV7 cortical spheroids just after implantation. e: Two-photon image of DIV7 cortical spheroid axons just after implantation. f: Two-photon image of DIV7 cortical spheroid axons just after implantation. g: Two-photon image of the same cortical spheroids as in *d* five days after implantation. h: Two-photon image of the same cortical spheroid axons as in *e* five days after implantation. i: Two-photon image of the same cortical spheroid axons as in *f* five days after implantation. j: Two-photon image of the seeding well of cortical spheroids 14 days after implantation. The green autofluorescent fibrous ingrowth is of unknown origin. k: Implantation track of the GelSH-GelNB conduit in the mouse brain. White arrows indicate the location of the remaining gelatin conduit. The conduit was fabricated shorter than the 3 mm required to reach the dLGN. Longer conduits tended to collapse during the PBS washout before they could be mechanically stabilized by transglutaminase crosslinking. Hoechst stain. dLGN: dorsolateral geniculate nucleus. l: Implantation track of the GelSH-GelNB conduit in the mouse brain. Red arrows indicate the location of the remaining collagen tube. Bright-field image. dLGN: dorsolateral geniculate nucleus. m: Cranial window and biohybrid implant with GelMA conduit 22 days after implantation. White arrows indicate the location of the collagen tube.

### Supplementary information

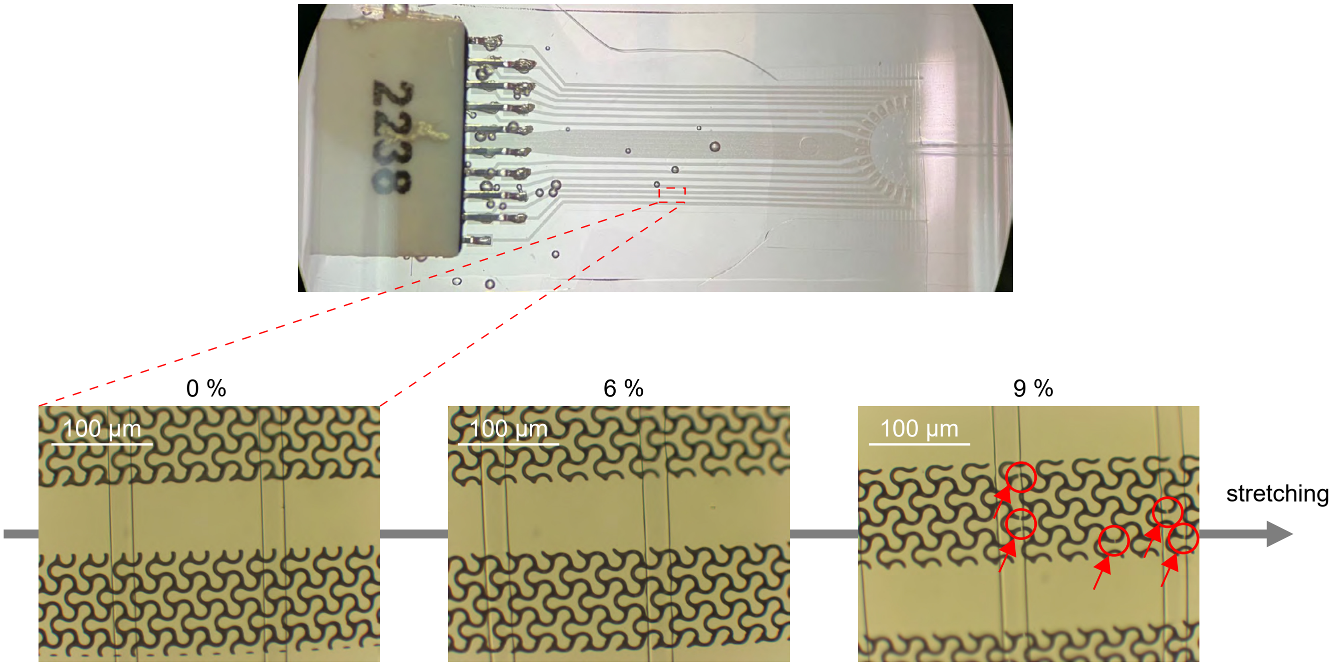


**Fig. S1**: **Uniaxial stretching of a microstructured track** Micropatterned tracks stretched under a microscope. The patterning confers mechanical compliance to the tracks, which start to crack at around 9% strain (the cracks are highlighted in red circles).

#### 6.2 Directional axon growth

To achieve functional independence of stimulation nodes and high bandwidth, axons were required to grow towards the implanted conduit and avoid converging onto neighboring nodes. To optimize unidirectional growth of axons, we focused on mechanical guidance motifs that were previously shown to achieve directional connectivity of up to 4 nodes [59]. Building on this work, we designed a set of 20 microstructures that systematically varied in specific features to identify the combination of motifs resulting in maximal bandwidth. The 20 designs (Supplementary Fig. S3) varied in features such as the channel width, the number and placement of 2-joints, the integration of so-called rescue loops, the specific design of 2-joints, the addition of spiky tracts, and finally the design of the final joining lane (see Supplementary Fig. S3c for illustration of motifs). These 20 designs systematically increased in complexity: whereas design 0 was treated as a negative control with no geometric features promoting directional joining, design 20 integrated all motifs potentially improving directed growth (Supplementary Fig. S3b).

##### 6.2.1 Axon tracking analysis

To characterize the directionality of our 20 microstructures at scale, we imaged axons over the initial stages of outgrowth from seeding nodes to the output channel using confocal microscopy. To assess growth direction of the hundreds of extending axons within our microstructures, we trained a convolutional neural network to detect growth cones in timelapse frames (Supplementary Fig. S4a). The multiple object detection model was adopted from the YOLOv3 architecture [60], with the modification of adding temporal context frames (past and future) to the input layer of the network. After linking axon identities through time using minimum cost flow optimization on a graph of detected axons within frames, we were able to calculate distances towards the output and neighboring nodes at single axon resolution throughout the lifetime of a detected growth cone (Supplementary Movie 7,8). The model reached a multiple object tracking (MOT) accuracy of 0.61, which measures the number of false positives, false negatives, and identity switches normalized to the number of ground truth labels. The model is published under the MIT license and publicly available at https://github.com/LoaloaF/axtrack.

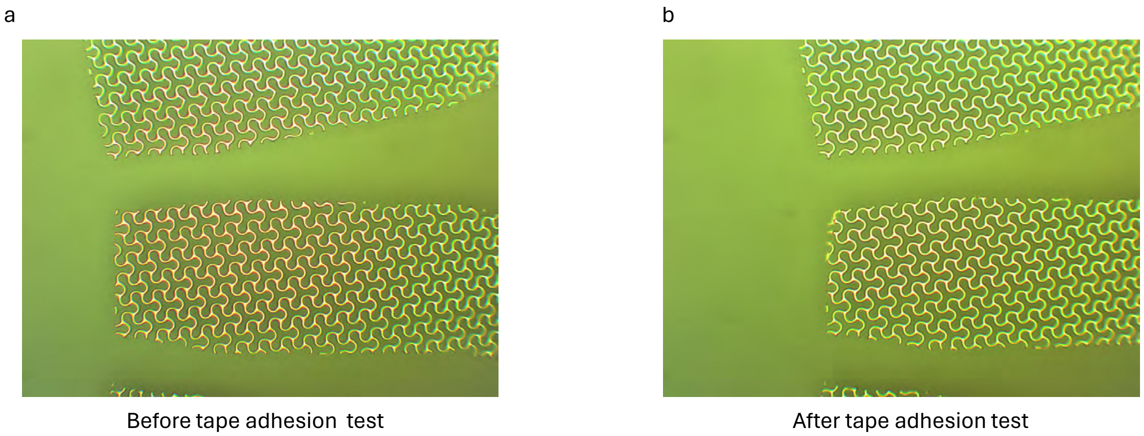


**Fig. S2**: **Tape adhesion test** Microscope pictures of a track on PDMS a: before and b: after the tape test.

Tracking the distance towards the output channel over time (Supplementary Fig. S4b) enabled us to read out the growth direction at single-axon resolution. Beyond directionality, the growth cone dislocation between frames allowed us to compare outgrowth velocity and the likelihood for axons to stagnate within the micro channels. We found that thinner channels (3 and 1.5 $\mu$m in designs 3 and, 4 respectively) exhibited significantly faster outgrowth and reduced growth stagnation (Supplementary Fig. S4c,f)). Features such as the addition of 2-joints and rescue loops had a weak negative effect on outgrowth velocity (Supplementary Fig. S4d,e). The effect of different features on directionality is shown in Supplementary Fig. S4g-l. One significant design feature was the number of 2-joints, or merging structures. Using one or three instead of none yielded significantly lower backwards growth, whilst not affecting the forward bias. Conversely, forward bias was significantly impacted by the placement of 2-joints but did not have an effect on backward growth counts. Joining channels early yielded a significant increase in forward growth of approximately 10%. Lastly, the number of rescue loops had an impact on both forward,- and backward growth. Implementing three instead of one or none rescue loops prior to each 2-joint reduced the proportion of backward-growing axons. Forward bias was significantly increased between three and fewer than three rescue loops. Finally, no significant effects were observed for angled versus straight rescue loop designs, spiky tracts, different variants of 2-joints, and modifications to the final lane design.

In conclusion, we identified a set of design motifs that had a significant effect on the directional growth of extending axons. In the final device used for *in vivo* implantation, additional modifications had to be introduced to accommodate the large stretchable electrode pads.

#### 6.3 Axon growth on PDMS

For the biohybrid implant we need axons to grow on PDMS. Initial trials showed poor axonal growth behaviour on PDL coated PMDS. Therefore, we tested the effect of different surface coatings and PDMS surface treatments on axonal growth within an axonal guidance structures mounted onto either glass (control) or PDMS. In all cases, coatings were forced into the axon guidance system by applying a vacuum (desiccation chamber for about 30min).

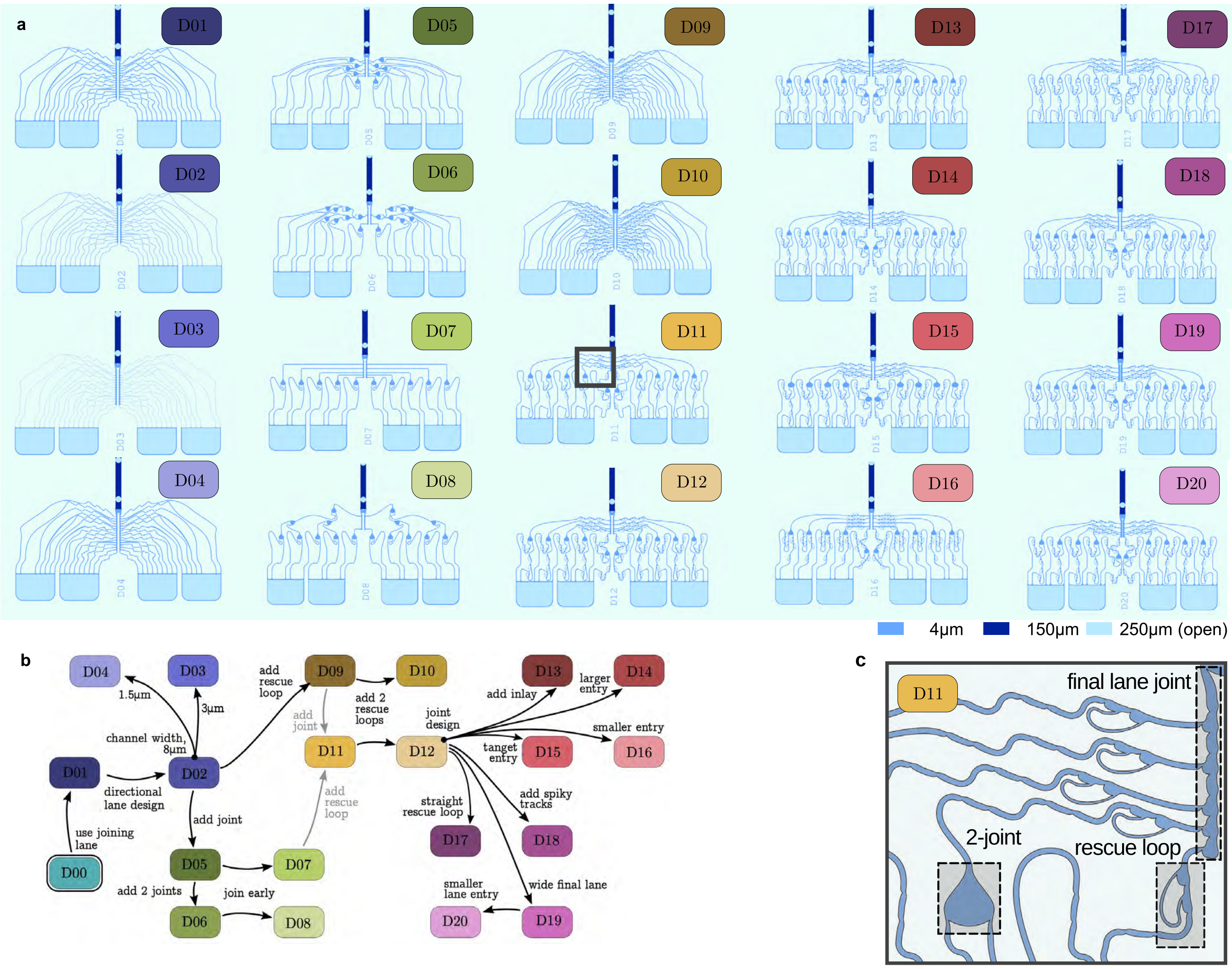


**Fig. S3**: **Exploring PDMS microstructure designs for enhanced unidirectional axon growth and merging** a: 20 microstructure designs implementing combinations of different design motifs. b: Overview of the design features distinguishing the different microstructures. Starting from Design 00 (D00), the arrows represent the implemented design feature changes. Blue designs vary in channel width (2-4), green designs integrate 2-joints (5-8), yellow designs add rescue loops (9-12), red designs (13-16) explore different 2-joints, and purple designs (17-20) vary features such as the final joining lane. c: Enlarged view of Design 11 illustrating specific features varied across designs.

To measure axon growth growth behavior we determined the fluorescence intensity of retinal axons over the channel length at different timepoints and counted how many channels that exited the seeding wells were filled with axons (Supplementary Fig. S6). As we suspected uncured monomer contamination to be one of the reasons for poor axonal growth, we tested how washing of the PDMS in Toluene for 24 h or postcuring at 80 °C would affect axonal growth under different surface coating conditions (Supplementary Fig. S6). Results showed that the postcuring heat treatment negatively affected axonal growth (Supplementary Fig. S6b) and the number of axon filled channels (Supplementary Fig. S6c). Both toluene wash and plasma treatment (30 s, maximum power, PDC-32G Harrick Plasma, Ithaca, NY, USA) resulted in a high fraction of axon filled channels that is comparable to the control condition on glass. The samples that were not pre-treated or toluene washed showed initially good axonal growth reaching the end of the 5 mm long channel, but started to retract after 22 days (Supplementary Fig. S6c). A stable axon bundle that persisted for more than 30 days without collapsing was achieved only in the plasma activated sample and control condition on glass.

##### 6.3.1 PDL coating

The bottom layer PDMS was coated with 2 mL of 0.05 mg/ml poly-D-lysine (PDL) solution (P7280, Sigma-Aldrich) and incubated at 4 °C overnight. The PDL solution was washed 3x with PBS and once with sterile DI water and dried for 1 h in the hood.

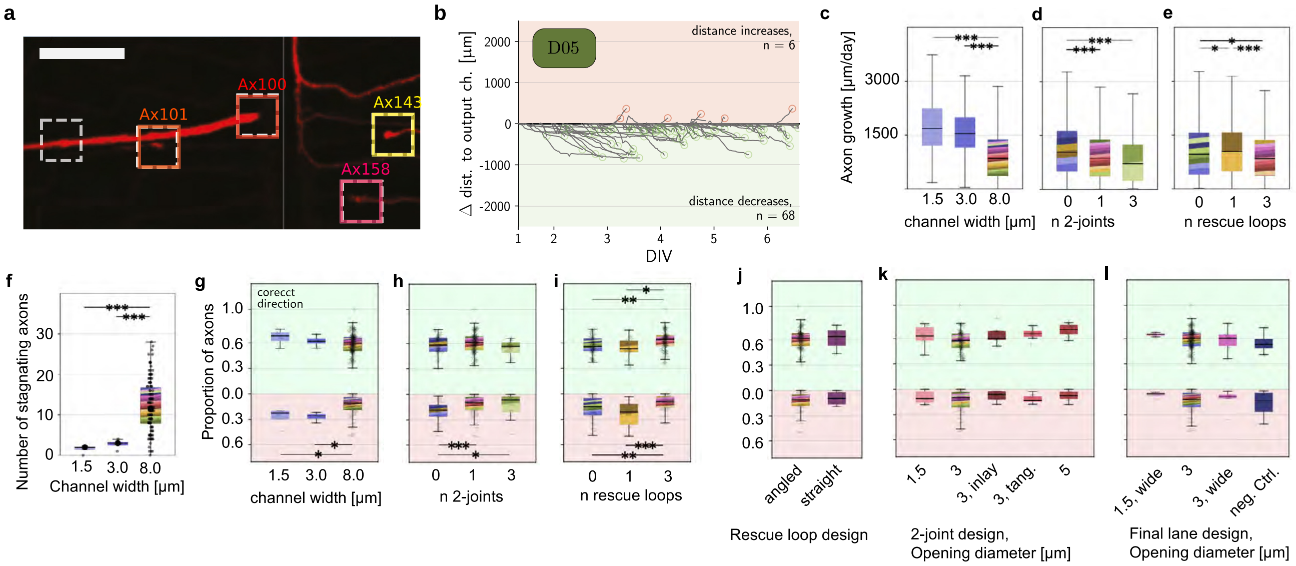


**Fig. S4**: **Time lapse of extending growth cones linking design features with unidirectional axon growth** a: Representative growth cone detection example in time lapse recording. Dashed boxes are predicted, colored ones are ground truth. Scale bar = 90 $\mu$m. Spheroids of 3000 retinal neurons/spheroid were seeded manually into the microstructure. Time-lapse recordings were started about 24 h after seeding the spheroids and performed on a confocal laser scanning microscope (CLSM) (FLUOVIEW FV3000, Olympus) using a 20x objective (UPLFLN20XPH, NA=0.5), 800 $\mu$m pinhole, a gain of 580-600 V, and laser power of 0.8-1 %. Timelapse recordings were captured at a resolution of 0.49 x 0.62 $\mu$m and an inter-frame-interval of 25 min. b: Inferring directionality from distance-to-output channel exemplified in design 5. Each of the gray lines represents an axon identity. Subtracting the initial distance to the output channel yields delta distances. Green or red circles mark axons that grew at least 50 $\mu$m in the correct or incorrect direction, respectively. Count is given in corners. c: Axonal growth speed depending on the channel width. Box plots with 1.5 x inter quartile range (IQR). Colors represent the 20 designs composing the distribution according to the color code in Supplementary Fig. S3b. d: Axon growth speed depending on the number of channel merges (joints) until the nerve channel. e: Axon growth speed depending on the number of rescue loops that redirect axons into the correct direction. f: Number of axons that stop moving along their path (stagnating axons). g-l: Proportion of correctly (green),- and incorrectly (red) growing axons in one half of a PDMS micro structure depending on different microstructure features. As indicated by the circles in panel b, only axons with delta distances above 50 $\mu$m were included to calculate proportions. Kruskal-Wallis test was used for non-parametric group comparisons, subsequently, the single comparisons were made using Mann-Whitney-U test with Holm-Bonferroni correction. * indicates $p<0.05$, ** $p<0.005$, and *** $p<0.0005$.

##### 6.3.2 Laminin coating

50 $\mu$L laminin (stock concentration: 1 mg/mL, 11243217001, Sigma-Aldrich) stock was thawed on ice and diluted in 5 ml Neurobasal plus medium (A3582901, Gibco) for a final concentration of 10 $\mu$g/mL. For experimental groups involving laminin coating, the surface of the dish was covered by a 2 mL laminin solution and incubated overnight at 37 °C in the incubator, or at 4 °C for 48 h. Finally, laminin solution was washed 1x with PBS and 2x with sterile DI water to avoid formation of salt crystals. DI water was immediately replaced with medium.

##### 6.3.3 Coating by desiccation

PDMS structures were mounted onto PDMS using 1:40 hexane diluted PDMS and cured at 80 °C for 2 h. After curing, PDL (0.05 mg/ml) was added directly (3 mL) and desiccated into the structures. Once the channels were filled, PDL was incubated for 1 hr at RT followed by 3 PBS washes with 10 minutes waiting between every wash. Finally, structures were washed with water, laminin (3 mL) was added and incubated overnight in the incubator. The next day, laminin was washed once with PBS (10 min) and 3x with sterile DI water (10 min). Medium was immediately after desiccated into the microstructure.

#### 6.4 Effect of PDMS monomer extraction on axonal growth

Our goal was to analyze how PDMS monomer extraction using ethylacetate would affect axonal growth and how it compares to plasma treated PDMS surfaces. For the screen, we cut out small circular PDMS pieces from

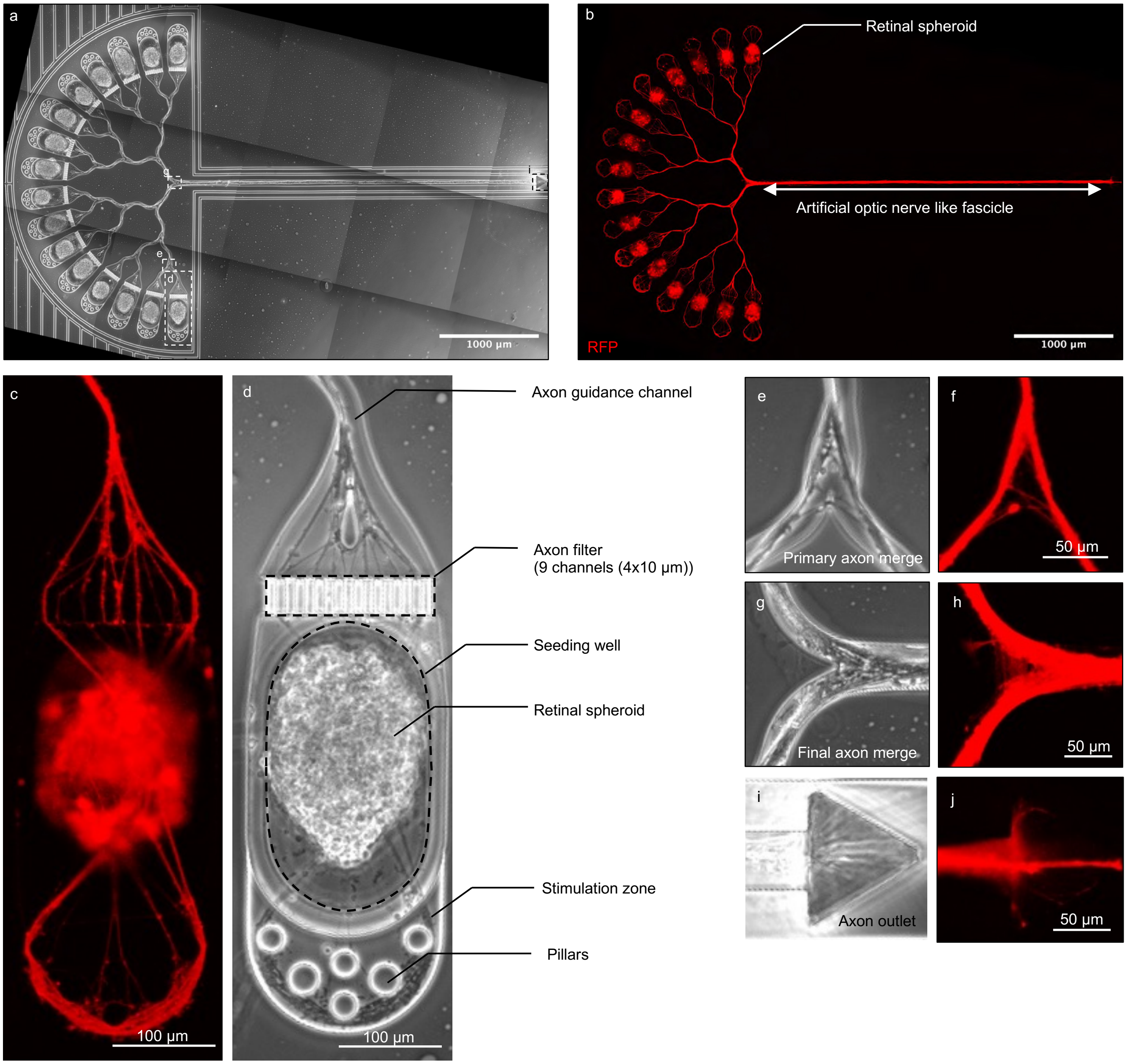


**Fig. S5**: **Axon growth in PDMS microstructure mounted on glass** a: Biohybrid microstructure mounted on glass. b: Fluorescence image of axon growth in biohybrid structure at DIV7. c: Retinal spheroid and axon growth behaviour in the seeding wells d: Illustration of the different features of the seeding well. e: Primary merge structure in which axons from 2 seeding wells merge towards the nerve channel. f: Fluorescence image of axons merging. g: Final merge structure that directs all axons into the nerve channel. h: Fluorescence image of axons merging. i: Axonal outlet at the end of the nerve channel. The triangular shape favours axons migrating up and out of the channel instead of turning back into the channel. j: Fluorescence image of the axons growing out of the nerve channel.

extracted and/or plasma activated PDMS sheets (produced by mixing PDMS monomer with crosslinker at a 1:10 ratio and cured at 80°C over night) using a 3 mm biopsy puncher and placed them into a 48-well plate. 30k rat dissociated primary cortical neurons from E18.5 embryos were seeded into each well. The results of our first set of experiments showed that axons and neural somata showed very poor adhesion to the non-extracted PDMS surface (Supplementary Fig. S7a,b). PDMS coated with PDA, PDL and laminin showed the highest neuronal coverage. Surface plasma activation (2 min, maximum power PDC-32G Harrick Plasma, Ithaca, NY, USA) resulted in a neuronal surface coverage that was indistinguishable from the control polystyrene surface independent of the coating condition (Supplementary Fig. S7c,d). Interestingly when repeating the experiment with a new batch of PDMS and neurons we did not observe any difference between extracted or non-extracted PDMS anymore. We concluded that PDMS monomer extraction or plasma activation should be performed for all implants to reduce the high variability

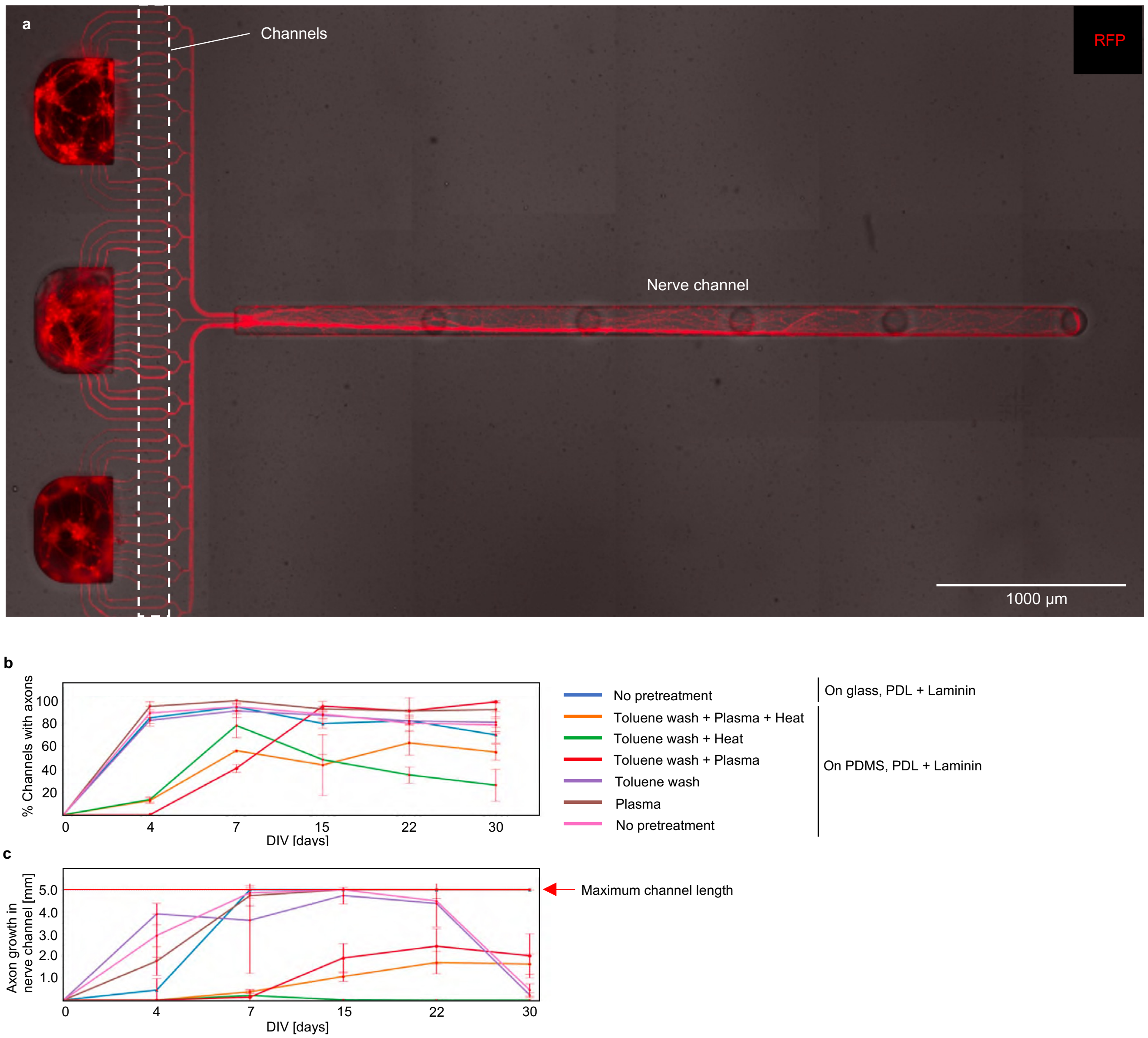


**Fig. S6**: **Effect of different PDMS coatings on axon growth** a: Fluorescence image of a first prototype of the biohybrid structure used to assess axonal growth behaviour. b: Quantification of the fraction of channels filled with axons depending on different surface treatments, error bars are SD, n=3. c: Quantification of the axon growth over time inside the nerve channel depending on different surface treatments, error bars are SD, n=3.

in axonal growth on PDMS.

#### 6.5 PDMS extraction

PDMS was extracted using ethyl acetate on a roller mixer (3x exchange every 6-8 h). Structures were washed 2x in 2-propanol for at least 30 min and subsequently dried.

#### 6.6 In situ fabrication of GelSH-NB conduits onto the PDMS microstructure

The collagen conduits used in Supplementary Fig. S12 were too big for implantations into mice and required a tedious manual alignment and gluing procedure. Therefore our goal was to replace the PDMS axon guidance channel with a hydrogel conduit that can be directly fabricated on top of the axon outlet of the PDMS microstructure. First, we tuned the geometric parameters to achieve a conduit with an outer diameter of about 300 µm and an inner diameter of about 150 µm. A cuvette was filled with a mixture of UV-crosslinkable GelSH-NB gel and exposed to the DMD projection pattern. The excess gel was washed out with PBS (1x) and conduits were post-crosslinked with transglutaminase (5U/ml) to increase mechanical stiffness (Supplementary Fig. S9a-b). For better visibility,

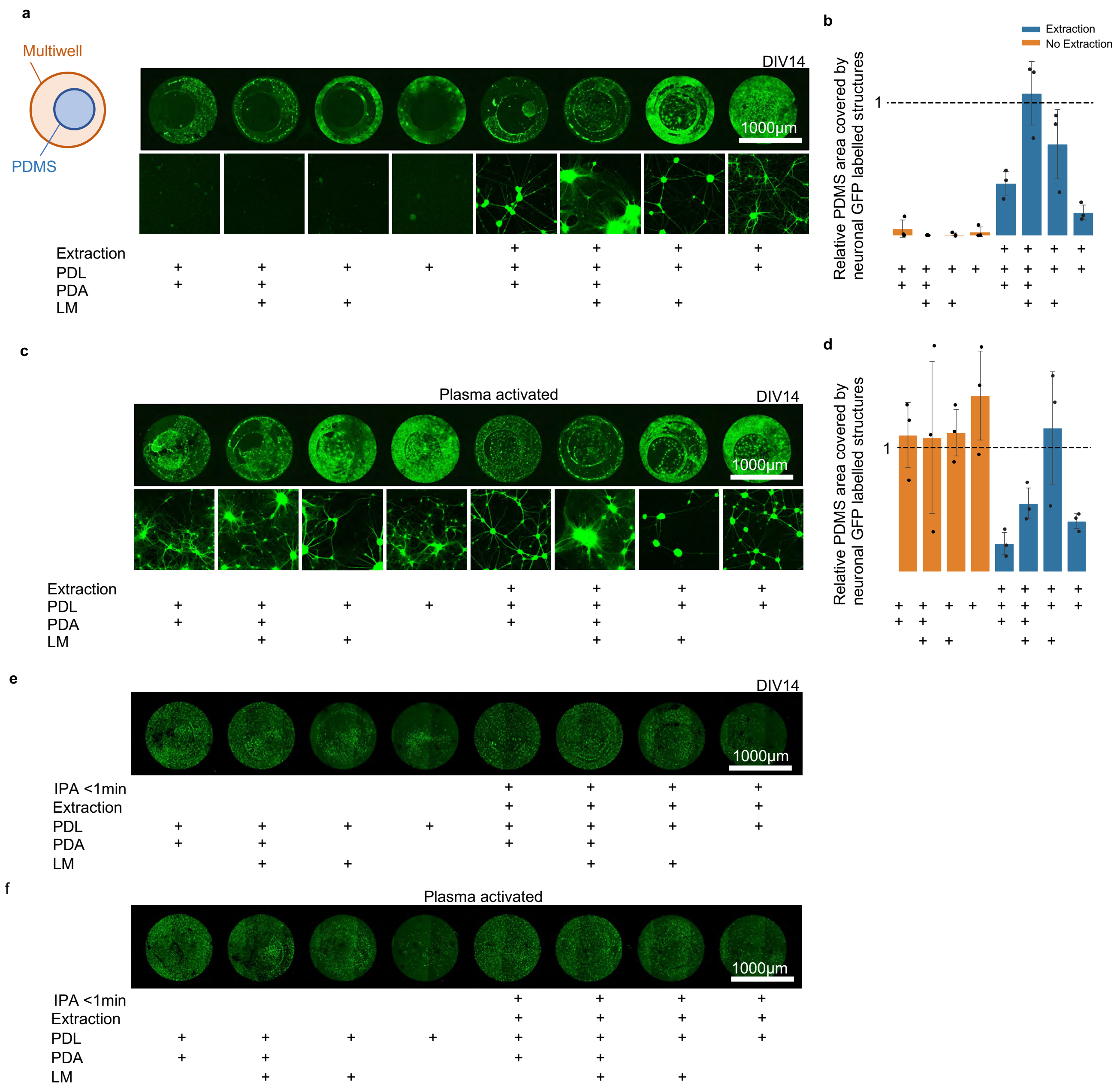


**Fig. S7**: **Effect of PDMS ethylacetate extraction on axon growth.** a: Relative PDMS surface area covered with GFP labelled neuronal structures (somata, axons and dendrites) compared to the control surface (Polystyrene) depending on extraction and surface coatings analyzed at DIV14. b: Quantification of data shown in a. c: PDMS surface area covered with GFP labelled neuronal structures (somata, axons and dendrites) relative to the control surface (Polystyrene) depending on extraction and surface coatings analyzed at DIV14. d: Quantification of data shown in c. e/f: Second iteration of the experiment indicating no effect of extraction illustrating the high variability without PDMS extraction. Neurons were labelled with a GFP expressing AAV at DIV7. To determine area coverage of GFP+ neurons, a defined threshold value (0.95 %) was applied to all images for binarizing. PDL = 0.1 mg/ml, Polydopamine (PDA) = 2 mg/ml, Laminin = 50 µg/ml, Plasma activation for 2 min, n=3 duplicates per condition. Error bar, standard deviation

the conduits were labelled with Rhodamine. We confirmed a hollow conduit structure by inserting an acupuncture needle (Supplementary Fig. S9b) and using confocal imaging (Supplementary Fig. S9b',b"). We next aimed to fabricate the conduits directly onto the axon outlet of the axon guidance structure. For alignment, GelSH-GelNB conduits were first fabricated onto the Ibidi dish without the implant (Supplementary Fig. S9c). We marked the location of the conduits and then removed them. We next aligned the implant axon outlet onto the marked location and fabricated the conduits directly onto the axon guidance microstructure according to the procedure described

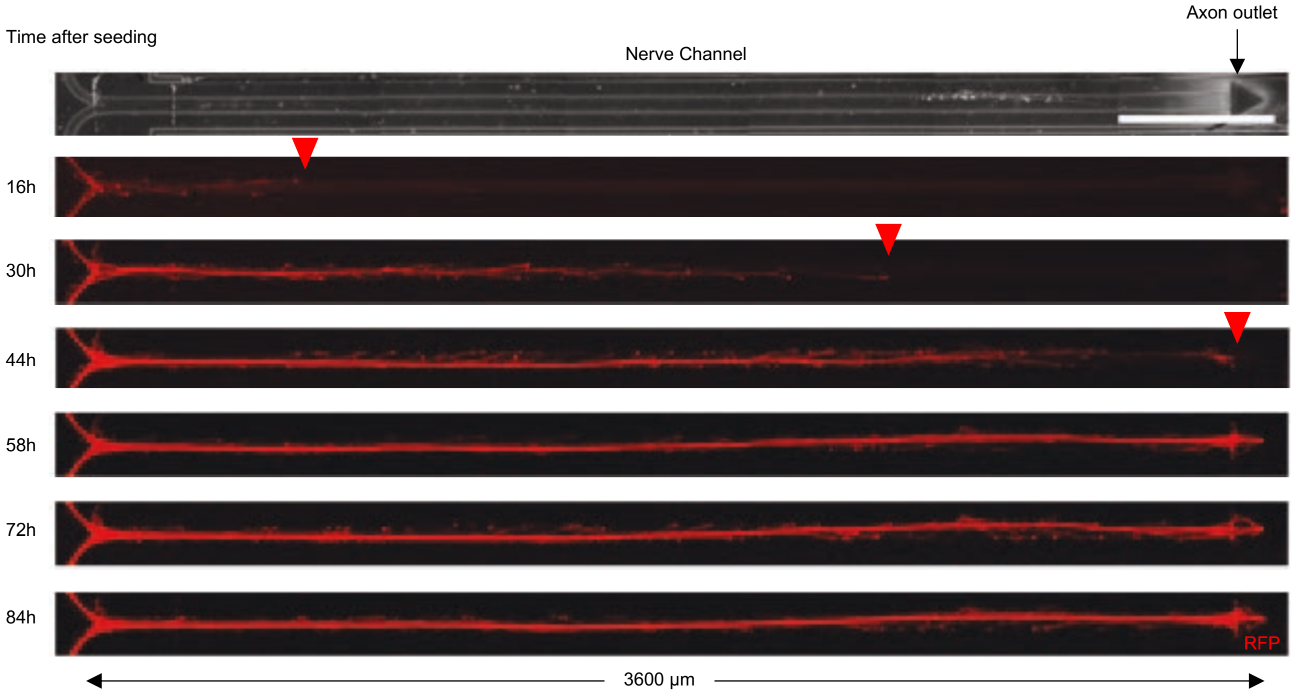


**Fig. S8**: **Axon growth in PDMS nerve channel** Snapshots of a timelapse movie to illustrate the speed of axonal growth and nerve like structure formation within the PDMS nerve channel. First axons reach the end of the 3.6 mm long nerve channel after 44 h.

before (Supplementary Fig. S9d,e). The height of the conduit can be determined by the amount of medium filled into the well.

We enhanced the hydrophilicity of PDMS microfluidic devices through plasma activation, which significantly improved the adhesion between the GelSH-GelNB conduits and the PDMS surfaces. This follows our previous findings that photocrosslinked GelSH-GelNB hydrogels exhibit better adherence to hydrophilic materials like glass that contain Silanol (Si-OH) groups which enhance the hydrogel bonding and ionic interactions with water-based materials such as hydrogels [54]. In the present case, plasma activation introduces polar groups (hydroxyl, carboxyl, carbonyl groups, etc.) and increases the surface roughness of the PDMS. This increases the hydrogen bonding between the GelSH-GelNB resin and the PDMS. It's important to note that these interactions might exist even before crosslinking; however, in their uncrosslinked state, the hydrogel is too fluid and conformal, lacking the necessary force and structural integrity to adhere effectively. Here, photocrosslinking also increases the crosslinking density and network rigidity of the hydrogel, which can enhance adhesive interactions between the hydrogel and plasma-activated PDMS. Furthermore, photocrosslinking also causes network shrinkage and expels water from the gels. A lower water content could reduce the lubricating effect between the hydrogel and glass, allowing for stronger direct interactions between the polymer chains of the hydrogel and the PDMS surface.

#### 6.7 FIB-SEM imaging of the axon bundle within the nerve channel

For FIB-SEM of the nerve grown *in vitro* within the PDMS microstructures, RGC spheroids of 800 cells/spheroid were seeded into the biohybrid structure and imaged before fixation at DIV7. The PDMS channel was cut open on the top and temporarily covered with a PDMS slab to enable fixation and embedding with the resin.

##### 6.7.1 Dehydration and critical point drying

Samples were fixed and poststained in osmium tetroxide and uranyl acetate as described above, except for treatment with thiocarbohydrazide. After dehydration in ethanol the samples were then critical point dried from ethanol with liquid $CO_2$ using a critical point dryer (CPD-931, Tousimis, USA). Samples were mounted and vacuum sputter coated as described above, and imaged using a scanning electron microscope (Merlin, Zeiss). Images were acquired using an InLens SE-detector with an acceleration voltage of 2 kV, 100 pA at a working distance of 5 mm.

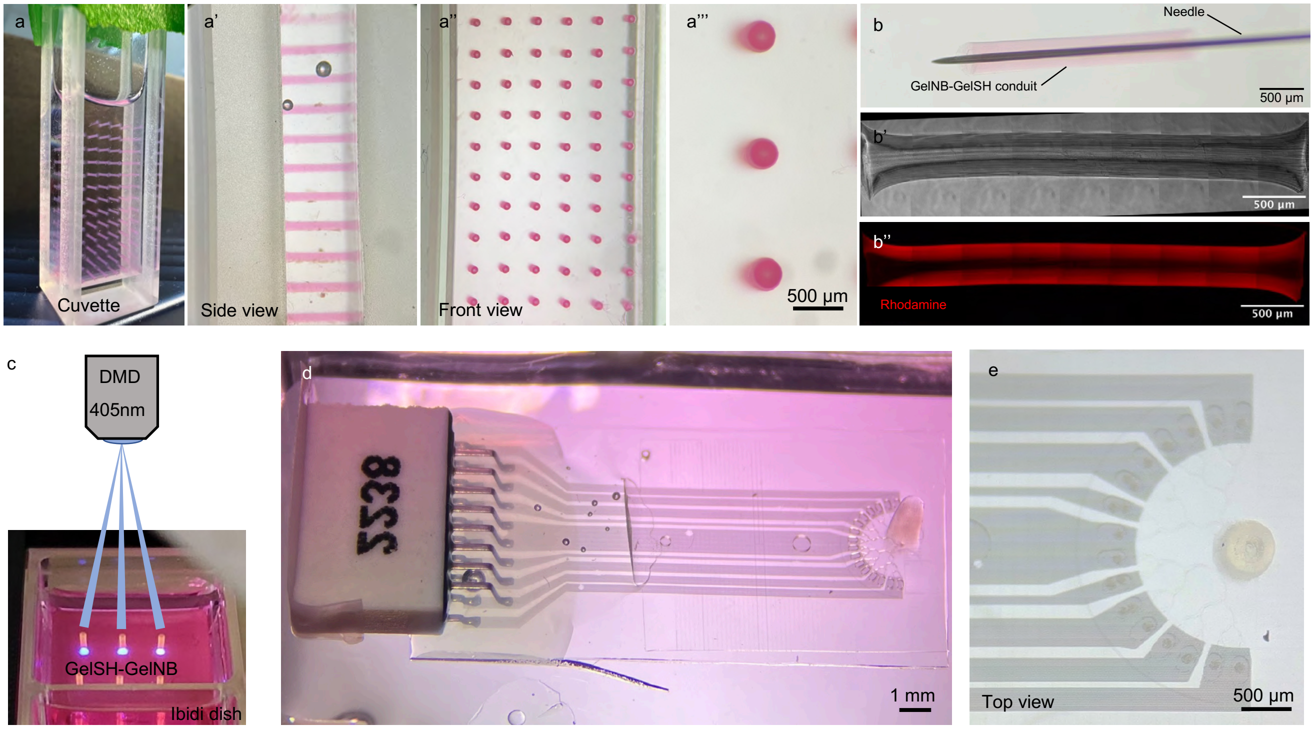


**Fig. S9**: **GelSH-NB conduit fabrication** a: Array of GelSH-GelNB fabricated conduits containing Rhodamine. a': Side view of the hydrogel conduits. a": Front view of the hydrogel conduits. a"': Zoom in of a". b: GelSH-GelNB conduit with an acupuncture needle inside. b': GelSH-GelNB conduit. b": Fluorescence image of the GelSH-GelNB conduit. c: Image illustrating how the conduits are fabricated. d: Biohybrid implant with a GelSH-GelNB conduit. e: Top view of the biohybrid implant with the hydrogel conduit fabricated on top of the axon outlet.

##### 6.7.2 Chemical fixation, dehydration and resin embedding

Samples were fixed in 2.5 % glutaraldehyde (EM grade; Polysciences Europe GmbH, Hirschberg an der Bergstrasse, Germany) and 2 % formaldehyde (EM grade; Polysciences), in 0.15 M Na-cacodylate buffer supplemented with 2 mM $CaCl_2$ and transported to the imaging facility ScopeM. The PDMS microstructure was removed and samples were overlaid with fresh fixative. After washing in cacodylate buffer (0.15 M, pH 7.2), cells were postfixed in 1.5 % potassium ferrocyanide (Merck) and 2 % osmium tetroxide (Polysciences), supplemented with 2 mM $CaCl_2$, followed by 1 % thiocarbohydrazide (Sigma), 2 % osmium tetroxide alone, and 1 % aqueous uranyl acetate (Polysciences). Samples were then dehydrated in a series of solutions with increasing ethanol concentrations (25 %, 50 %, 70 %, 90 %, 100 %) and finally infiltrated with Epon resin (Fluka, Buchs, Switzerland). All steps until resin infiltration were performed in a BioWave Pro+ Tissue Processor (Ted Pella Inc., Redding CA, USA). For en bloc embedding, the mold of the culture dish was filled with resin and after polymerization the glass substrate, on which the cells were grown, was removed to expose the cells with the basal side facing upwards. For thin-layer plastification, the culture dishes were set at an angle of approx. 80° and excess resin covering the cells was allowed to drain off before polymerization. That way, the cells can be analyzed with the apical side facing upwards. Samples were mounted onto SEM-stubs (Plano GmbH, Wetzlar, Germany) using a silver-filled epoxy glue (Epo-Tek H20E; Electron Microscopy Sciences, Hatfield PA, USA), and vacuum sputter coated with 7 nm Pt-Pd using a CCU-010 (Safematic GmbH, Switzerland). Cross-sections were prepared in a FIB-SEM (Helios 5UX, Thermo Fisher Scientific) and imaged using an ICD detector at an acceleration voltage of 2 kV, 0.1 nA.

#### 6.8 Axon growth into glass conduits

Here our goal was to see how axons transit from the axon guidance structure into a hydrogel filled glass conduit and how axons behave inside the tube. We used an acupuncture needle to align and glue the glass conduit onto a PDMS seeding mask using UV-curable PDMS. After complete curing, the needle was removed and the seeding mask with the attached tube was aligned onto the axon outlet of the axon guidance structure. Next the tube and plasma activated PDMS microstructure were filled with matrigel by applying a drop of matrigel diluted in NB medium (1:1) onto the top of the conduit and seeded with cortical spheroids (Supplementary Fig. S11a,b). Within 32 days axons transit into and reach the top of the about 3 mm long glass conduits (Supplementary Fig. S11c,d,e,f).

#### 6.9 Axon growth inside collagen conduits

Here we asked (1) how we can mount a collagen conduit that is used for peripheral nerve repair onto the PDMS axon guidance structures (2) how axons will transit into the conduit and (3) how they will extend into a target matrigel structure. Therefore the collagen conduit was mounted on top of a PDMS microstructure with an exit hole and glued onto it using UV-curable PDMS (Supplementary Fig. S12b,c). Next the conduit was aligned and glued onto the exit triangle of the axon guidance structure and filled with a 50% matrigel solution. Within 12 days axons transited from the PDMS microstructure into the collagen conduit and distributed within the conduit lumen (Supplementary Fig. S12d,e). We finally inverted the microstructure and inserted them into a PDMS microwell filled with matrigel to simulate an implantation *in vitro* (Supplementary Fig. S12a). Within 2 days after the *in vitro* implantation axons transited into the target matrigel (Supplementary Fig. S12d,f).

##### 6.9.1 Fabrication of the collagen conduit

The study outlines the development of collagen-based nerve guidance conduits (NGCs) using spinning mandrel technology, following the method previously established [61]. The NGC's primary support structure consisted of insoluble collagen (2.5 %, w/w) swollen in 1 M acetic acid and homogenized with a high-speed mixer at 10,000 rpm (Polytron®, Kinematica, Lucerne, Switzerland) for approximately 1 min. The researchers procured Microfibrillar Collagen Hemostat (Avitene) from BARD (Oberrieden, Switzerland). The collagen dispersion was evenly applied onto a spinning mandrel coated with gold (diameter: 1.5 mm) using a syringe. The solvent was then dried under laminar air flow overnight, resulting in a structurally stable device. The resulting tubes were neutralized by incubating them in 0.1 M di-sodium hydrogen phosphate (pH 7.4) for 1 hour and eventually cut into 14 mm long specimens. The collagen NGC specimens underwent a dehydro-thermal treatment (DHT) at 110 °C and 20 mbar for 5 days, which cross-linked them through physical means. Resulting NGCs measured in the range of 1.5 mm inner diameter and 2.5 mm outer diameter in the hydrated state. This method has been demonstrated to produce NGCs with exceptional mechanical properties and biocompatibility, as shown in our previous studies [62, 61, 63].

| Implantation | Spheroid age [days] | Cell type | Implant type | Implant location | Max. days with fluorescent neurons | Max. days with fluorescent axons |
|---|---|---|---|---|---|---|
| 1 | DIV0 | Cortex | PDMS channel | Epidural | 23 | 5 |
| 2 | DIV0 | Cortex | PDMS channel | Epidural | 5 | 5 |
| 3 | DIV0 | Cortex | PDMS channel | Epidural | 5 | 0 |
| 4 | DIV0 | Cortex | PDMS channel | Epidural | 5 | 0 |
| 5 | DIV1 | Retina | PDMS channel | Epidural | 4 | 4 |
| 6 | DIV1 | Retina | PDMS channel | Subdural | 8 | 4 |
| 7 | DIV1 | Retina | PDMS channel | Epidural | 8 | 4 |
| 8 | DIV1 | Cortex | PDMS channel | Epidural | 4 | 4 |
| 9 | DIV1 | Cortex | PDMS channel | Epidural | 0 | 0 |
| 10 | DIV7 | Retina | PDMS channel | Epidural | 9 | 0 |
| 11 | DIV7 | Retina | PDMS channel | Epidural | 9 | 5 |
| 12 | DIV7 | Retina | PDMS channel | Epidural | 5 | 0 |
| 13 | DIV7 | Cortex | PDMS channel | Epidural | 5 | 5 |
| 14 | DIV7 | Cortex | PDMS channel | Epidural | 7 | 0 |
| 15 | DIV1 | Cortex | GelMA conduit | Epidural | 4 | 4 |
| 16 | DIV1 | Cortex | GelMA conduit | Epidural | 16 | 6 |

Table S1: *In vivo* experiments summary

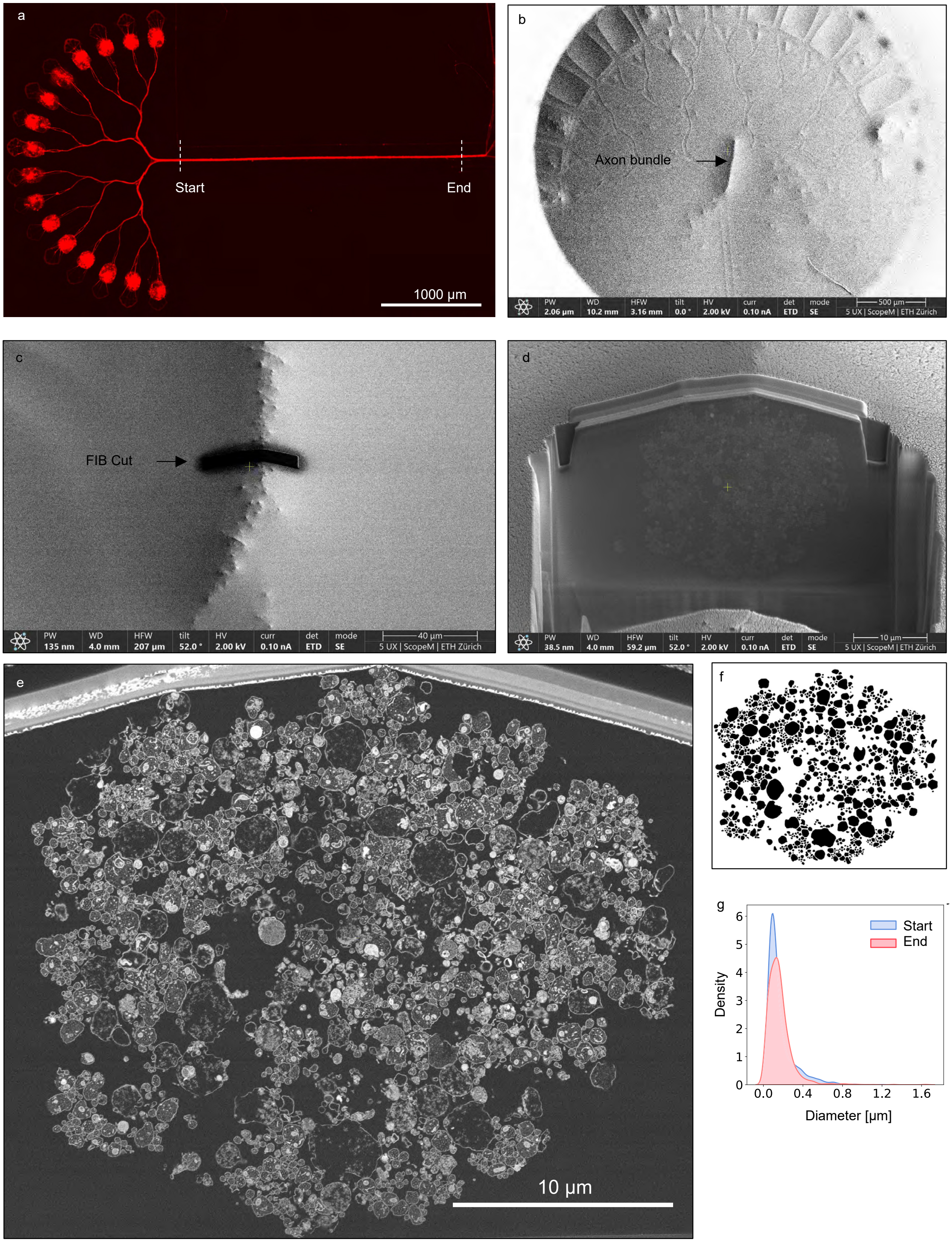


**Fig. S10**: **Focused ion beam (FIB) imaging of the axon bundle** a: Fluorescence image of the biohybrid structure mounted on glass used for FIB-SEM imaging. b: SEM image after the fixation procedure. The axon bundle got dislocated during the fixation procedure. c: FIB cut into the axon bundle. d: Magnification of the FIB cut into the axon bundle. e: SEM Image of the axon bundle cross section. f: Segmentation of the axon bundle for quantification in g. g: Kernel density distribution of the axon diameter distributions comparing cuts at the beginning of the axon bundle (start) and close to the outlet (end).

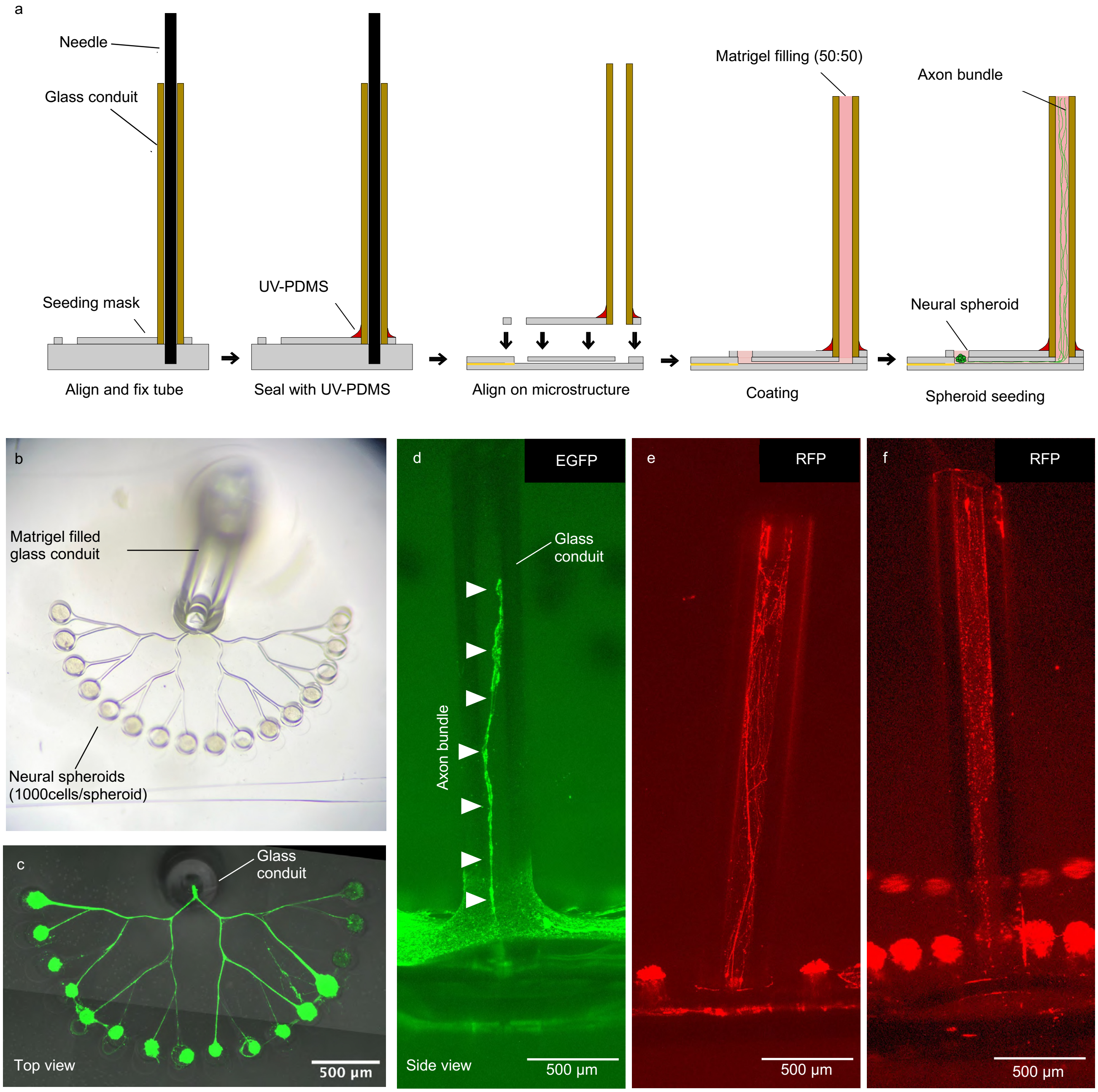


**Fig. S11**: **Axon growth within PDMS mounted glass conduits** a: Illustration of the fabrication procedure to mount the glass conduit onto the axon outlet of the axon guidance structure. b: Photograph of the axon guidance structure with the glass conduit mounted onto the axon outlet. c: Fluorescence image illustrating axonal growth after spheroid seeding. d-f: Side view maximum projection illustrating how axons form a nerve like structure within the matrigel filled glass conduit.

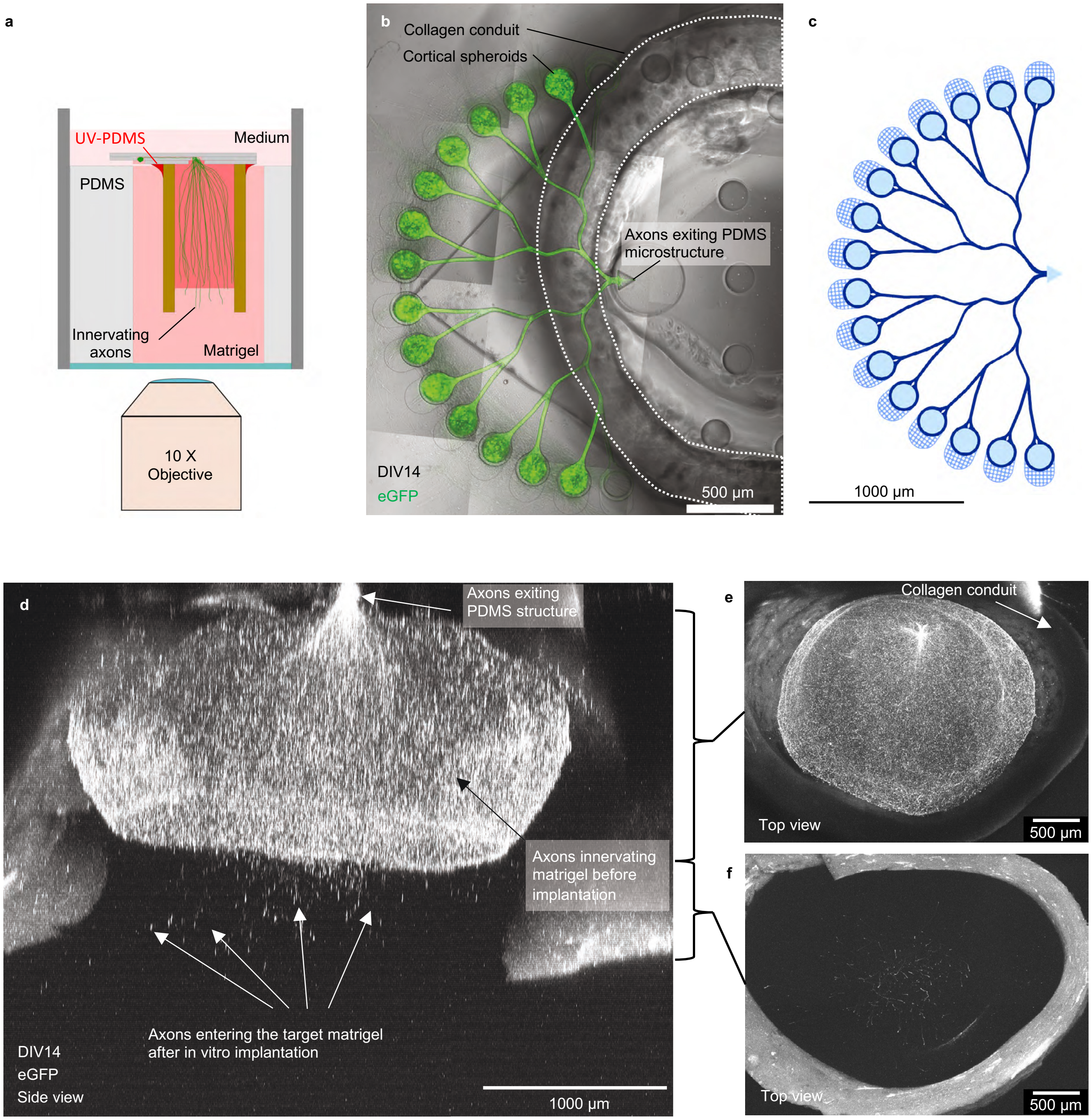


**Fig. S12**: **Axon growth within matrigel filled collagen tube** a: Schematic side view to illustrate how the spheroid containing biohybrid implant was inverted and injected into a 50:50 matrigel filled PDMS well. b: Top view of the matrigel filled collagen conduit mounted onto the axon outlet of the axon guidance structure. c: CAD layout of the used axon guidance structure. d: Side view maximum projection showing how axons exit the PDMS microstructure and grow into the matrigel filled lumen of the collagen conduit. After implantation axons continued to grow to infiltrate the target matrigel (White arrows). e: Top view maximum projection of the collagen tube showing axons covering the lumen of the conduit. f: Top view maximum projection of the upper part of the collagen tube illustrating axons innervating the target matrigel.

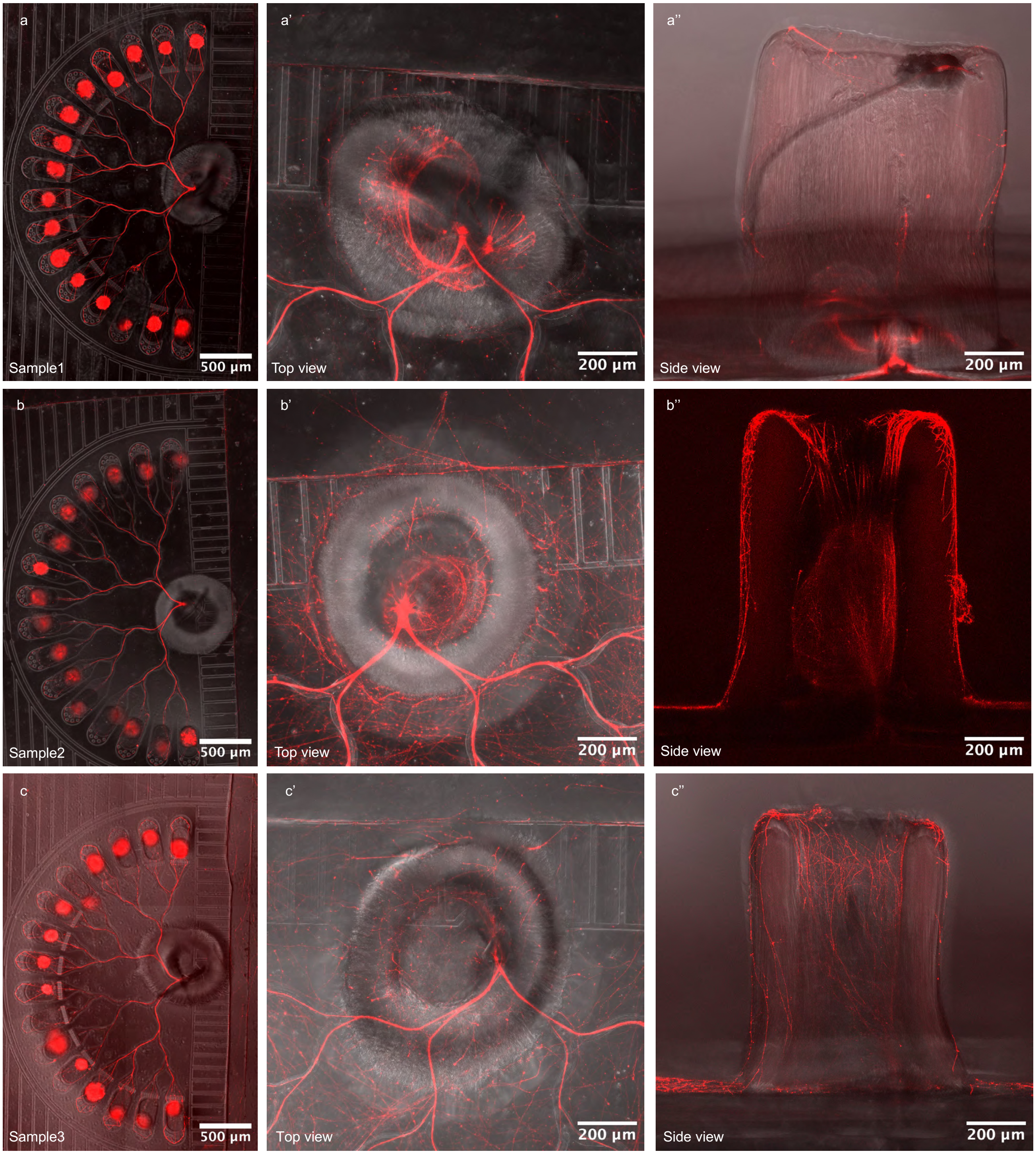


**Fig. S13**: **Axon growth within the GelSH-GelNB hydrogel conduits** a-c: Top view of axonal growth in the biohybrid structure with a fabricated GelSH-GelNB conduit of different samples. a'-c': Top view zoom in of the hydrogel conduit of different samples. a''-c'': Side view maximum projection of the hydrogel conduits to illustrate axonal growth from the PDMS microstructure into the conduit.
